# Heterochromatin-based silencing of a foreign tandem repeat in *Drosophila melanogaster* shows unusual biochemistry and temperature sensitivity

**DOI:** 10.1101/2025.07.31.667933

**Authors:** Tingting Gu, Elena Gracheva, Michael Lee, Wilson Leung, Sophia Bieser, Alixandria Nielsen, Adam T. Smiley, Nhi N.T. Vuong, Matthias Walther, Gunter Reuter, Sarah C. R. Elgin, Andrew M Arsham

**Author notes:** Baylor College of Medicine, Houston, TX 77030. St. Jude Children’s Research Hospital, Memphis, TN 38105.

## Abstract

Eukaryotic genomes are packaged into chromatin, a regulatory nucleoprotein assembly. Establishment, maintenance, and interconversion of chromatin states is required for correct patterns of gene expression, genome integrity, and survival. Transcriptionally repressive heterochromatin minimizes mobilization of transposable elements and limits expansion of other repetitive DNA, but mechanisms for recognition of the latter sequences are not well established. We previously demonstrated in *Drosophila melanogaster* that transcripts derived from *1360* and *Invader4* transposon insertions can trigger local conversion of transcriptionally permissive euchromatin to heterochromatin through the piRNA system, but only in a subset of genomic locations near existing blocks of heterochromatin. Here we show that a ~9 kb tandem array of the 36-nucleotide *lac* operator (*lacO*) sequence of *Escherichia coli* can form ectopic heterochromatin at a similar subset of sites, resulting in variegating expression of an adjacent reporter gene. Heterochromatin Protein 1a (HP1a) and histone deacetylation are required for *lacO* repeat-induced silencing, but, contrasting with previously described Position Effect Variegation (PEV), we do not observe increased histone H3 lysine 9 methylation. Silencing is effective at 25°C and suppressed at 18°C (in contrast to canonical PEV, which is enhanced at 18°C), indicating involvement of a temperature-sensitive component. Temperature switching experiments show that *lacO* repeat-induced heterochromatin formation is reversible throughout larval development following an HP1a-dependent initiation step in the early embryo. We conclude that the *Drosophila* nucleus can recognize a completely foreign tandem repeat as a target for heterochromatin formation, and that the heterochromatin structure established is distinct from that of endogenous tandem arrays.

## Introduction

Eukaryotic genomic DNA is packaged in chromatin, a dense, dynamic, and heterogeneous nucleoprotein complex. Chromatin packaging and the three-dimensional organization of the nucleus impact gene activity, establishing patterns that enable the selective expression required for cell differentiation and development in multicellular organisms. Chromatin also helps define the boundaries of key genome structures, including telomeres and centromeres, which transmit epigenetic information over cellular, organismal, and evolutionary time scales (reviewed by (Janssen et al. 2018)).

The primary repeating unit of chromatin is the nucleosome, with a core of 147 bp of DNA spooled around a barrel-shaped histone protein octamer. The chromatin state is generally defined by the post translational modification of the N-terminal tails of histone proteins (Strahl and Allis 2000) and the differentially associated chromosomal proteins (reviewed by (Elgin and Reuter 2013)). Chromatin can broadly be categorized into two types: euchromatin – gene-rich, loosely packed, and transcriptionally accessible – is characterized by high levels of histone acetylation; in contrast, heterochromatin – gene-poor and repeat-rich, more densely packed, and generally transcriptionally inactive – is characterized by low histone acetylation, distinct histone methylation, and the presence of heterochromatin proteins such as Heterochromatin Protein 1a (HP1a). “Heterochromatin” is a useful oversimplification of a large and complex portion of the genome. Heterochromatic domains in fact comprise many subdomains (some with active genes) containing different kinds and lengths of repetitive sequences and transposable elements (TE) with diverse histone marks. Heterochromatin accumulates these repetitive sequences in part by mitigating their deleterious impact through inhibition of transcription and recombination. In addition to the major blocks of heterochromatin that define centromeres and telomeres, short heterochromatic domains can also be found interspersed throughout otherwise euchromatic regions. Some of these regions may be characterized by TEs or other repeats (reviewed in (Janssen et al. 2018)).

Because heterochromatin silences genes it is imperative that the cell correctly identifies regions of the genome to target for heterochromatin assembly. To explore this issue, we earlier established a transgene system in *D. melanogaster* whereby the addition of a repetitious sequence from the *1360/Hoppel* DNA transposon or from the *Invader4* retrotransposon can tip the balance, converting euchromatin to heterochromatin. The presence of the TE sequence stochastically silences an adjacent reporter generating Position Effect Variegation (PEV), a well-known indicator of heterochromatin packaging. Genetic and biochemical analyses have shown that in this case, the change is dependent on the piRNA system (Sentmanat and Elgin 2012).

piRNA-mediated silencing of TEs by heterochromatin has been investigated in detail by numerous labs (reviewed by (Czech et al. 2018)), but much less is known about the silencing of other deleterious repeats like tandem arrays of short sequences. Could tandem repeats also drive ectopic heterochromatin formation in this reporter system? If so, would small RNA or other sequence-specific signals be involved, or is there something broadly inherent to DNA tandem repeats that can trigger silencing? Cloning and manipulating repetitive DNA fragments is difficult, so to begin to address this question we used an existing a 256-copy array of 36 bases of prokaryotic DNA (from the *lac* operon (*lac*O) of *E. coli*), using a P element vector to insert this fragment into the *Drosophila* genome. Here we show that this *lacO* array can induce ectopic heterochromatin formation in a subset of genomic regions. These regions substantially overlap those that we previously found susceptible to TE-induced heterochromatin formation (Haynes et al. 2006; Huisinga et al. 2016). However, unlike common TEs, the *lacO* repeat sequence used here can be defined as “foreign”, *i*.*e*. not previously seen by the organism. It is not present in the *D. melanogaster* reference genome (Release 6 plus ISO1 MT assembly) nor in a recent PacBio HiFi assembly which includes the highly repetitive regions of the *D. melanogaster* genome (Shukla et al. 2024). Thus the experiments reported here argue against recognition based on pre-existing genomic sequences (such as an RNAi mechanism) and instead point to properties inherent to repetitive DNA.

The heterochromatin formed at *lacO* repeats has a number of interesting and useful properties, with a distinct histone modification pattern and an unusual temperature sensitivity which we have exploited to probe the silencing process. We anticipate that the system we have developed will be useful in future studies examining how organisms defend themselves against invading non-TE repetitious sequences.

## Results

### A tandem array of repetitive foreign DNA directs HP1a-dependent silencing

Euchromatic genes placed in close proximity to heterochromatin by rearrangement or transposition can be stochastically silenced by heterochromatin spreading to produce a variegating phenotype. In particular, a *white* (*w*) transgenic reporter inserted into heterochromatin by *P* element mobilization can assume a PEV eye phenotype; the transgene appears to be packaged in the same fashion as nearby heterochromatic genes (Riddle et al. 2008; Riddle et al. 2012; Messina et al. 2023). We have exploited this phenomenon to ask whether an array of foreign tandem repeats can induce ectopic heterochromatin formation, using an adjacent *white* gene as the reporter.

Prior work from our lab has shown that the DNA transposon remnant *1360* can trigger HP1a-dependent silencing of an adjacent *hsp70-white* reporter gene, but only when inserted in specific genomic locations (Haynes et al. 2006; Sentmanat and Elgin 2012; Huisinga et al. 2016). In this investigation we used a *phiC31* landing-pad system to replace the *1360* sequence at insertion site 1198 (2L:20,094,149) with a ~9 kb tandem array of 256 copies of a 36 nucleotide *E. coli* lac operator sequence, *lacO* (Sasmor and Betz 1990; Li et al. 2003). The 1198 insertion site was identified using *1360* to screen for genomic sites where a variegating phenotype is dependent on the presence of a repetitious element (Sentmanat and Elgin 2012). Site 1198 is located in the 5’UTR of the *nesd* gene, within a gene-rich euchromatic region on the left arm of chromosome 2. This euchromatic location, while several megabases away from pericentric heterochromatin, is approximately 3 kb from an H3K9me2- and HP1a-enriched heterochromatin domain (Kharchenko et al. 2011) and about 10 kb from a piRNA cluster (Brennecke et al. 2007).

The insertion of 256 copies of the *lacO* sequence resulted in striking silencing of the *white* transgenic PEV reporter (Figure 1, top right), stronger than that observed with *1360* (Sentmanat and Elgin 2012). This silencing does not affect the expression of a nearby *yellow* transgene, consistent with prior reports showing that the strong *yellow* promoter is resistant to heterochromatin-induced silencing (Yan et al. 2002). We used flanking FRT (Flippase Recognition Target) sites to excise the *lacO* array in the germline by *in vivo* heat shock-induced FLP recombinase. This eliminated silencing, producing progeny with red eyes (Figure 1, bottom right) and confirming that the silencing is repeat-dependent.

**Figure 1.**
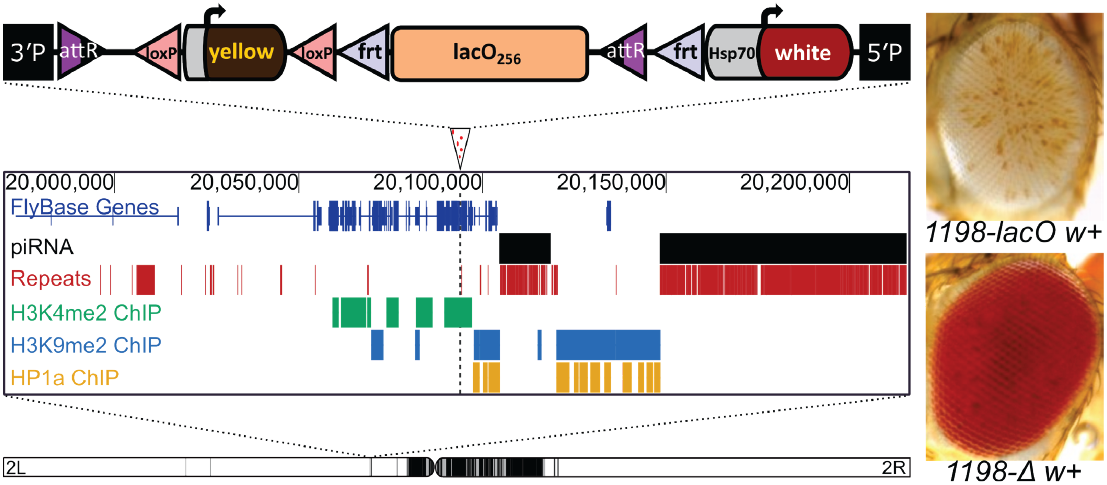
A tandem array of lacO repeats initiates variegated silencing of a reporter gene when inserted at site 1198. Top: schematic of the lacO hsp70-white reporter construct; middle: genome browser showing the lacO construct insertion site (marked by an arrowhead with dotted line below); the presence in the surrounding 250kb of piRNA clusters and repetitive DNA; and H3K4me2, H3K9me2, and HP1a maps derived from modENCODE ChIP data from S2 cell lines (Kharchenko et al. 2011). Bottom: position of the 1198 insertion site on the left arm of D. melanogaster chromosome 2 (Sentmanat and Elgin 2012). Black domains indicate pericentric heterochromatin; white domains indicate euchromatin. Inset: eye phenotypes in the presence (1198-lacO w+) and absence (1198-Δ w+) of lacO repeats in the construct.

In *D. melanogaster*, local heterochromatin formation is typically associated with high levels of Heterochromatin Protein 1a (HP1a, encoded by the gene *Su(var)205*). Consistent with a role in *lacO* repeat-induced silencing, variegation is significantly suppressed (Figure 2A) in flies heterozygous for *Su(var)205*^*5*^ (a frameshift truncating the protein after amino acid 10 (Eissenberg et al. 1992)) or *Su(var)205*^*2*^ (a point mutation disabling the chromo domain(Platero et al. 1995)).

**Figure 2.**
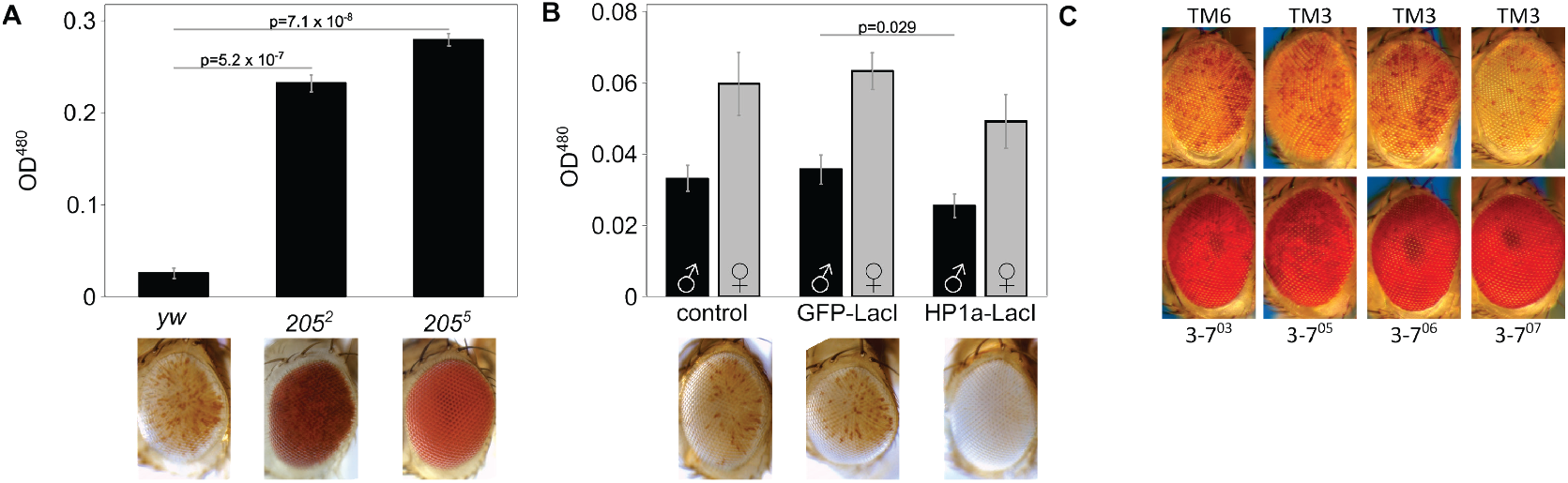
LacO repeat-induced silencing is HP1a-dependent. (A) HP1a mutations suppress variegation. 1198-lacO/CyO flies were crossed with one of two distinct HP1a mutant lines (Su(var)205^2^ or Su(var)205^5^) or a yw control. Variegation was suppressed in flies heterozygous for HP1a compared to controls. All pigment assays were performed on male flies, and all photographs are of male fly eyes except for the yw control. (B) Tethering of an HP1a-LacI fusion protein to the lacO array enhances silencing. 1198-lacO/CyO flies were crossed with mutants expressing heat shock-activated HP1a-LacI or GFP-LacI fusion proteins that bind to the lacO repeat array. HP1a-LacI recruitment resulted in significant loss of expression of the downstream hsp70-w reporter relative to yw and the GFP-LacI controls. Both males and females were tested in the pigment assay (black bars = males; grey bars = females); males are pictured. Two-tailed P-values comparing at least three replicates of the two groups at either end of the horizontal bar are shown in (A) and (B). P values were corrected for multiple tests; error bars show standard deviation. (C) Four heterozygous mutations in the heterochromatin-associated protein Su(var)3-7 suppress lacO repeat-induced silencing (males are pictured; mutation present, bottom row; controls, top row).

Fusion proteins with the *lacO*-binding domain of the *E. coli* LacI protein can be used to tether domains of interest to *lacO* DNA arrays (Memedula and Belmont 2003). Expression of a GFP-LacI fusion protein had no impact on reporter expression, indicating that protein binding *per se* at the 1198 site has no impact on local heterochromatin state. In contrast, expression of an HP1a-LacI fusion protein (Danzer and Wallrath 2004) enhanced variegation in the *1198-lacO* flies, providing supporting evidence that HP1a, and hence heterochromatin, play a role in *lacO* repeat-induced silencing (Figure 2B). Further confirming the involvement of HP1a, four distinct EMS-induced mutations in the gene for the HP1a-interacting partner *Su(var)3-7* (Cléard et al. 1997) are dominant suppressors of *lacO* repeat-induced variegation (Figure 2C).

### Silencing induced by the *lacO* repeat is insensitive to disruption of HP2 or of the Polycomb or RNAi complexes

Using PEV reporters, over 150 suppressors and enhancers of variegation (referred to respectively as *Su*(*var)* and *E(var*)) have been identified in *D. melanogaster* in genetic screens (reviewed in (Elgin and Reuter 2013)). Most of these encode proteins that are either components or post-translational modifiers of chromatin (Elgin and Reuter 2013). To further investigate the nature of the *lacO* repeat-induced variegating phenotype, we crossed *1198-lacO* males to females carrying mutations in genes coding for heterochromatin associated proteins (identified as *Su(var)* genes), visually scoring progeny for dominant suppression or enhancement of the PEV phenotype. Mutations in *Su(var)2-HP2*, the gene encoding HP1a binding partner and suppressor of variegation HP2 (Shaffer et al. 2006), have no impact on *lacO* repeat-induced silencing (Table 1). The Polycomb system maintains silencing of key developmental regulatory genes by a chromatin-based complex that is distinct from HP1a-associated heterochromatin formation. Not surprisingly, mutations in the Polycomb complex gene *Enhancer of zeste* (*E(z)*) had no impact on *lacO* repeat-induced PEV (Table 1).

**Table 1.**
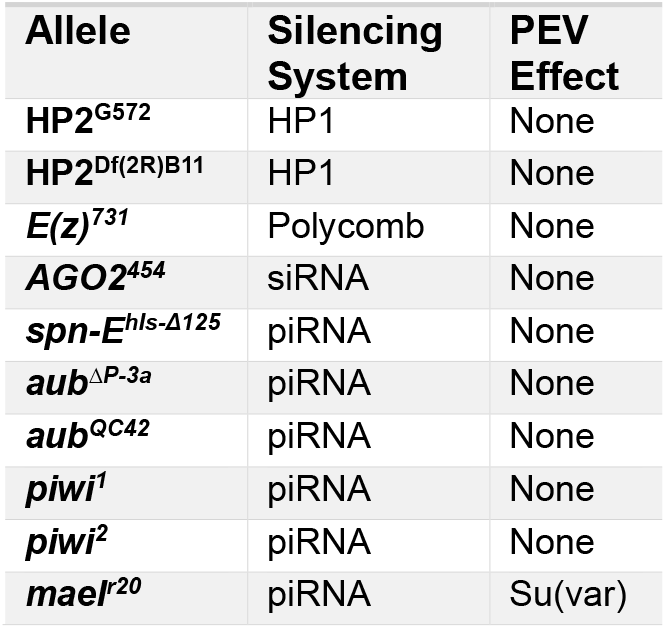
Repeat-induced silencing by lacO is not sensitive to mutations that disrupt HP2, or the Polycomb or RNAi complexes.

TEs in the *Drosophila* genome are targeted by the piRNA system for germ line silencing utilizing transcripts from endogenous piRNA cluster loci to provide target specificity (Czech et al. 2018). To evaluate the possibility that transcripts from the *lacO* repeats are activating endogenous RNAi-based genomic silencing systems, we used BLAST to compare a single copy of the 36 nucleotide *lacO* sequence and a 256-copy array of that sequence to two distinct *D. melanogaster* genome assemblies: the reference assembly (Release 6 plus ISO1 MT; NCBI RefSeq Assembly GCF_000001215.4), and a recent PacBio HiFi assembly that resolved the highly repetitive regions of the *D. melanogaster* genome (ASM4260644v1; NCBI GenBank Assembly GCA_042606445.1) (Shukla et al. 2024). No matches with an E-value < 0.05 were found, demonstrating the absence of *lacO-*matching sequences from existing piRNA clusters, as well as elsewhere in the genome.

Consistent with this, no loss of silencing was observed in flies with mutations in *Argonaute 2* (*AGO2), spindle E* (*spn-E*), *aubergine* (*aub*), and *piwi*, all known components of the piRNA system (Table 1 and Figure S3). This is in marked contrast to piRNA-dependent silencing of the TEs *1360* and *Invader4* inserted at the same genomic location (Sentmanat and Elgin 2012). A mutation in *maelstrom* (*mael*), a nucleic acid binding protein involved in piRNA regulation (Onishi et al. 2020) does suppress *lacO* PEV (Table 1); this effect was highly variable and only statistically significant in males. Given that mutations in other piRNA genes had no consistent impact on PEV, we suggest that *mael* facilitates *lacO* repeat-induced silencing via a mechanism unrelated to the piRNA system.

### Impact of developmental temperature on silencing by *lacO* repeats

The mottled pigmentation characteristic of PEV is thought to be established in two stages — a stringent and uniform initial silencing step during early embryogenesis followed by leaky maintenance. Stochastic decay of heterochromatin occurs during differentiation and development, allowing expression in some cells while maintaining silencing in others, giving the characteristic mottled appearance (Lu et al. 1996; Lu et al. 1998; Bughio et al. 2019; Bughio and Maggert 2022). While HP1a is required for both initial silencing and maintenance of heterochromatin, the piRNA machinery, required where the target element is a TE, is needed only for initial silencing (Gu and Elgin 2013). Prior studies of temperature dependence have found that PEV is enhanced by culturing at temperatures lower than that typical for the organism (Gowen and Gay 1933; Hartmann-Goldstein 1967). Thus we were surprised to observe a strong inverse temperature-dependence for *lacO* repeat-induced silencing, which was enhanced at 25°C and suppressed at 18°C.

To determine the developmental stage at which this difference is established, we cultured variegating *lacO* flies for several generations at the silencing-permissive temperature of 25°C and then carried out timed matings, shifting vials to 18°C at different stages in development. Transfers were done 24h after egg laying (‘embryo’); 48 hours after egg laying, when larval movement was visible on the food (‘L1’); 5 days after egg laying, when larval wandering was observed (‘L3’); and 7 days after egg laying, when most larvae had pupariated (‘pupal’). Control vials remained at 25°C for the duration of the experiment. Silencing was dramatically decreased in a developmental stage-dependent manner: the earlier the transfer, the more complete the suppression, indicating that *lacO* repeat-induced silencing requires active maintenance throughout embryonic and larval development (Figure 3A). No significant impact was seen when transfer occurred at the pupal stage, presumably because the pigment pattern had already been established.

**Figure 3.**
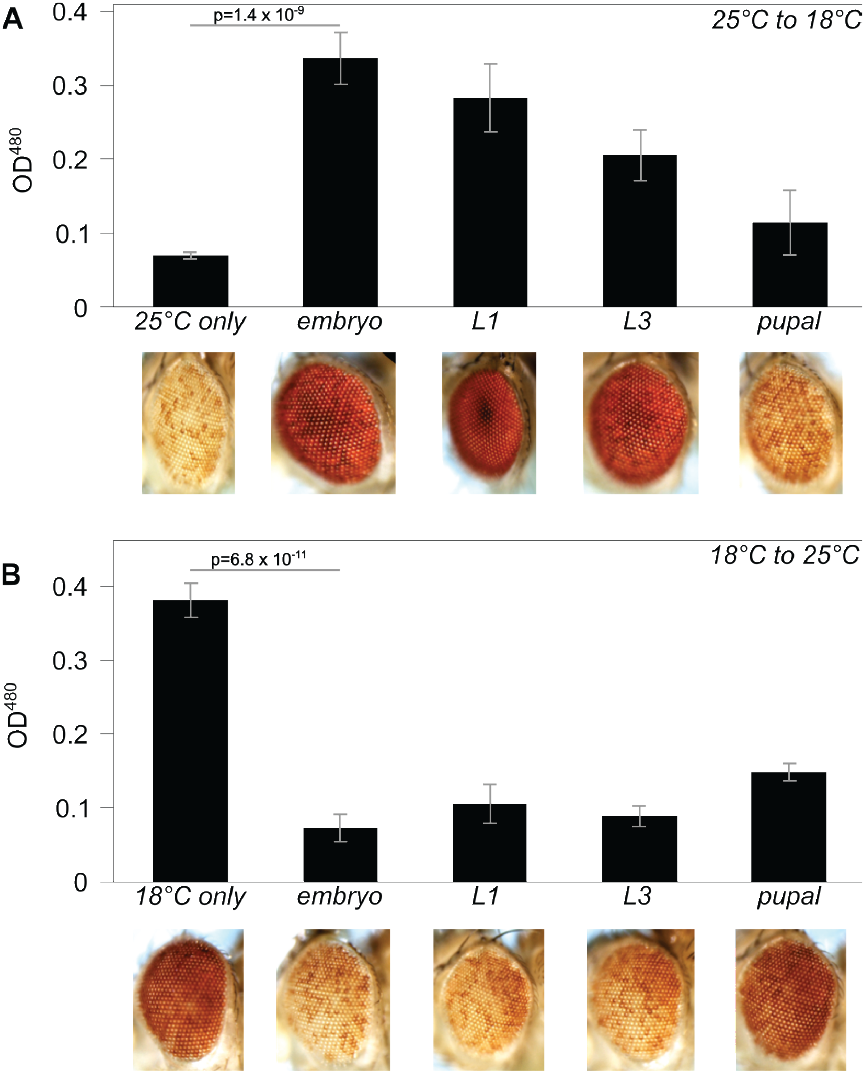
Impact of developmental temperature on silencing by lacO repeats. (A) Silencing triggered by the lacO repeat is robust when animals are raised at 25°C but is suppressed when animals are transferred to 18°C during embryonic or larval development. (B) Silencing triggered by the lacO repeat is weak in flies raised at 18°C but is enhanced when flies are transferred to 25°C during late embryonic or larval development. P values compare the averages of at least three vials maintained at the starting temperature to vials transferred during the embryo stage; error bars denote standard deviation. The partial effects seen on transfer at the pupal stage suggest that many of the flies in these samples had already completed their developmental use of the white product before the transfer, rendering the assay ineffective.

To test whether *lacO* repeat-induced heterochromatin could be assembled later in development, we reversed the temperature transition, culturing flies for several generations at 18°C before transfer of progeny to 25°C at different developmental stages, observed as described above (Figure 3B). Transfer to the silencing-permissive temperature any time before pupariation restored variegation. This suggests that the *lacO*-repeat site remains capable of initiating heterochromatin formation throughout development. The results imply the existence of a novel temperature-sensitive component required for maintenance of *lacO* repeat-induced heterochromatin, but not for initial recognition and licensing.

### Additional insertion sites support *lacO* repeat-dependent silencing

Initial testing of the impact of the *lacO* repeats was done at a single genomic location, 1198 (2L:20,094,149). To test the capacity of the *lacO* repeats to trigger silencing elsewhere in the genome we carried out a transposition screen, mobilizing the entire reporter P element with the *lacO* cassette intact (Figure 1, top) from the X chromosome and looking for *white* expression from new sites on the autosomes, as described previously (Sentmanat and Elgin 2012). The cassette includes a second reporter gene, the *yellow* (*y*) gene of *D. melanogaster* (scored by observing body pigmentation). The two reporter genes, *yellow* and *white*, provide additional insight into the establishment of ectopic heterochromatin. Several pilot screens were carried out to test feasibility and optimize workflows before scaling up. Out of 14,137 unique F2 progeny in the scaled-up screen, approximately 2% (288) had visible transgene expression. Of these, 82% (236 or 1.7% of all F2) displayed full reporter gene expression in eye and body color, suggesting insertions into euchromatic regions. These results demonstrate that in flies, tandem repeats do not trigger ectopic heterochromatin formation in most genomic locations. This is consistent with our prior findings using other repetitive DNA, specifically TE remnants (Haynes et al. 2006; Sentmanat and Elgin 2012; Huisinga et al. 2016).

About one fifth of the recovered mutants (52 or 0.4% of all F2) show some silencing of at least one of the two reporter genes, with the most common silencing phenotype (31 or 0.2% of all F2) being complete loss of *white* expression with visible expression of *yellow*. We also observed the converse: loss of *yellow* expression with strong expression of *white*. Hence in some cases, the expression of the two reporters is decoupled.

Approximately 0.06% (8 males out of 14,137) of all F2 in the scaled-up screen had variegated eye color; males from all three screens were used to establish new variegated stocks. We crossed the new lines to flies expressing the FLP recombinase in eye cells to assess whether removal of the *lacO* repeats early in eye imaginal disc development could reverse the silencing phenotype. To further characterize the silencing observed, we raised larvae from these lines at 18°C and at 25°C to look for impact of developmental temperature on silencing.

We used inverse PCR to map some locations of the new variegating insertions. Figure 4 shows the genomic location of new variegating lines in the context of our prior studies of repeat-triggered variegation using the *1360* transposon remnant (Sentmanat and Elgin 2012; Huisinga et al. 2016). The PEV phenotype in all five new *lacO* lines is dependent on the presence of the *lacO* repeats, as shown by the impact of excision of the *lacO* repeats with FLP. All inserts are also suppressed when larvae are raised at 18°C (Figure S4). The ability of the *lacO* repeats to establish ectopic heterochromatin in a context-dependent manner is thus not limited to a single location but is a general (albeit limited) *cis*-acting feature. Although there is no obvious pattern in the chromatin states of the different insertion sites (Table 2), many variegating insertions are in promoters or transcribed regions of euchromatic genes, presumably the best available target for insertion of such a large P element (Spradling et al. 1995). All five lines are within 8 Mb of a pericentric heterochromatin mass, but Lac018 is in the middle of the 2R long arm, generally considered a euchromatic domain. Analysis of flanking sequences (Table 2) reveals that all inserts except Lac018, including those within gene-rich regions, are located near >5 kb long blocks of repeats identified by RepeatMasker and Tandem Repeats Finder. No clear patterns in the distribution or type of such repetitive DNA is apparent. We also observed that the recovered *lacO* repeat-dependent silencing locations were often close to locations identified in previous studies looking at insertion sites of P constructs showing a *white* PEV phenotype dependent on an adjacent *1360* element (Sentmanat and Elgin 2012; Huisinga et al. 2016). Such regions, indicated by large clusters of light blue flags in Figure 4, might be considered “silencing prone.” In particular, a disproportionate number of variegating insertions map to short segments near the pericentric heterochromatin of chromosome 2.

**Table 2.**
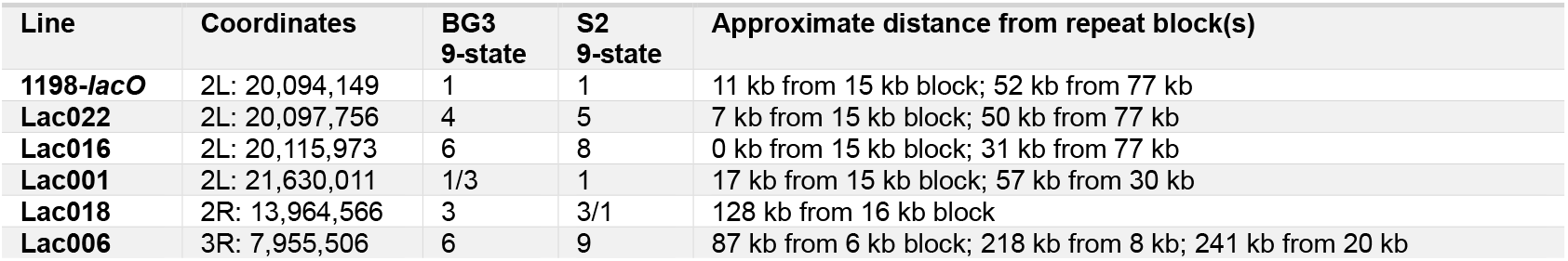
Characterization of newly isolated P element reporters exhibiting lacO repeat-induced white variegation, including cassette insertion coordinates, temperature-dependence, modENCODE chromatin states in BG3 and S2 cells, and proximity to blocks of repetitive DNA larger than 5kb.

| Line | Coordinates | BG3<br>9-state | S2<br>9-state | Approximate distance from repeat block(s) |
| --- | --- | --- | --- | --- |
| **1198-lacO** | 2L: 20,094,149 | 1 | 1 | 11 kb from 15 kb block; 52 kb from 77 kb |
| **Lac022** | 2L: 20,097,756 | 4 | 5 | 7 kb from 15 kb block; 50 kb from 77 kb |
| **Lac016** | 2L: 20,115,973 | 6 | 8 | 0 kb from 15 kb block; 31 kb from 77 kb |
| **Lac001** | 2L: 21,630,011 | 1/3 | 1 | 17 kb from 15 kb block; 57 kb from 30 kb |
| **Lac018** | 2R: 13,964,566 | 3 | 3/1 | 128 kb from 16 kb block |
| **Lac006** | 3R: 7,955,506 | 6 | 9 | 87 kb from 6 kb block; 218 kb from 8 kb; 241 kb from 20 kb |

**Table 3.**
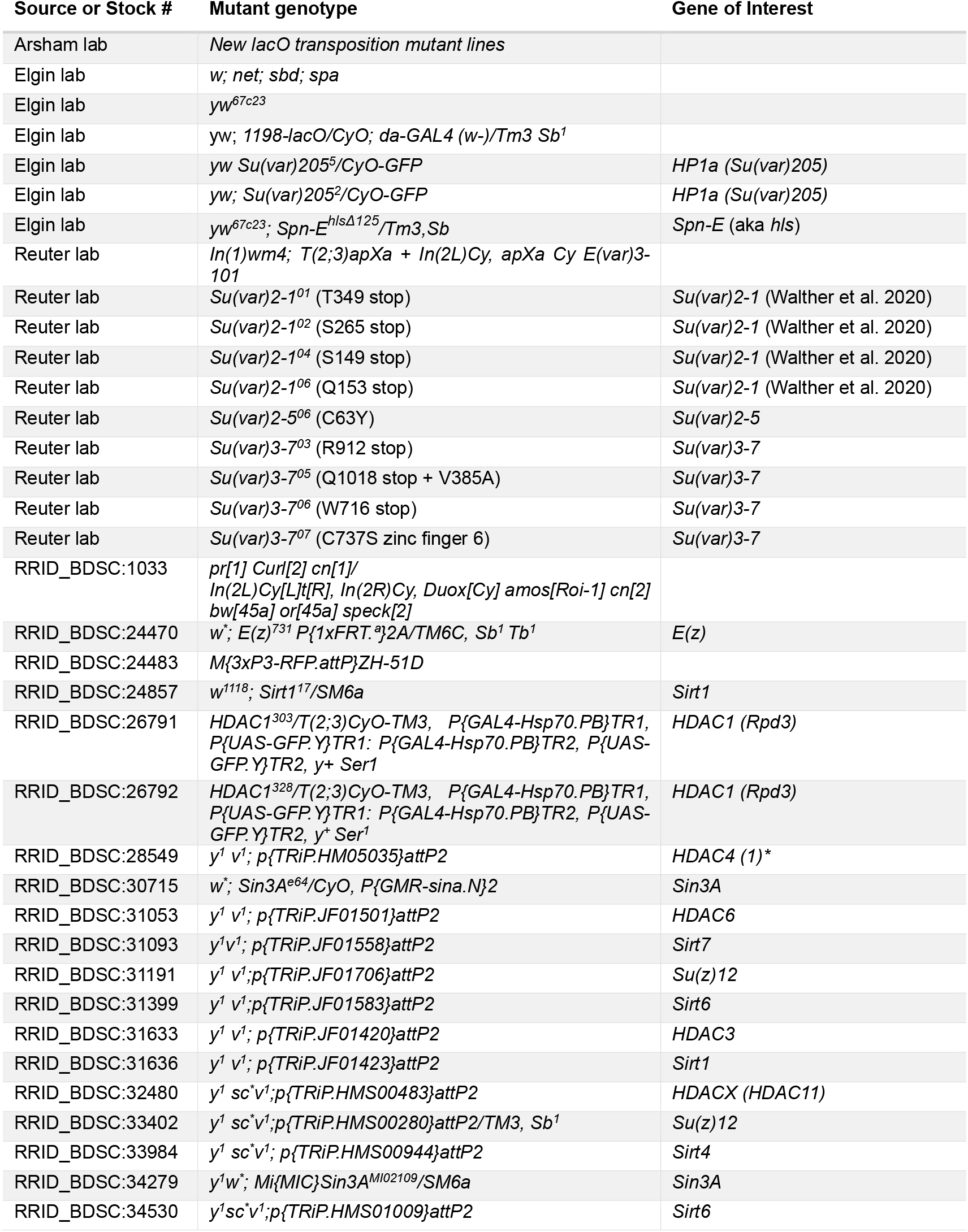
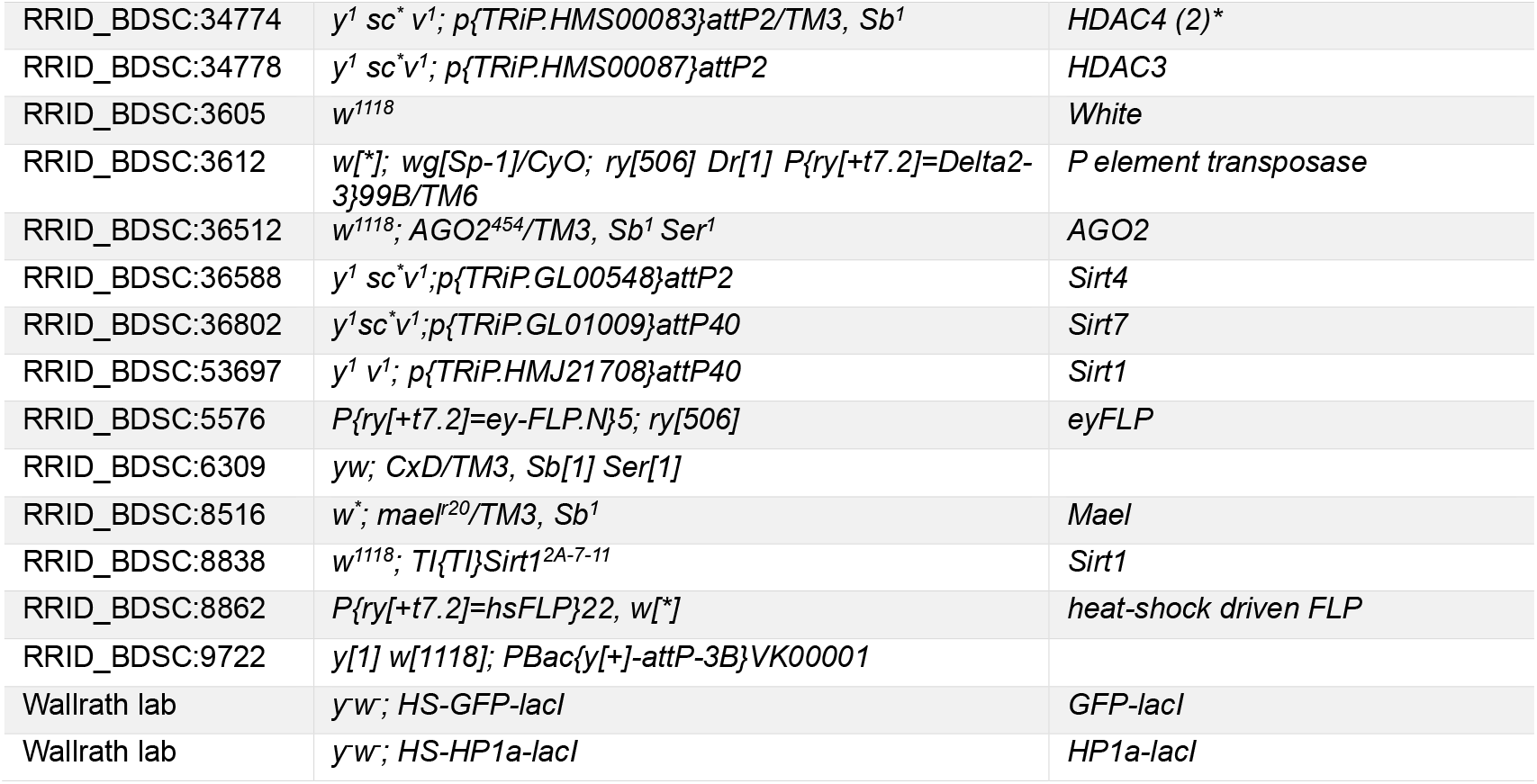
Drosophila Stocks used.

**Figure 4.**
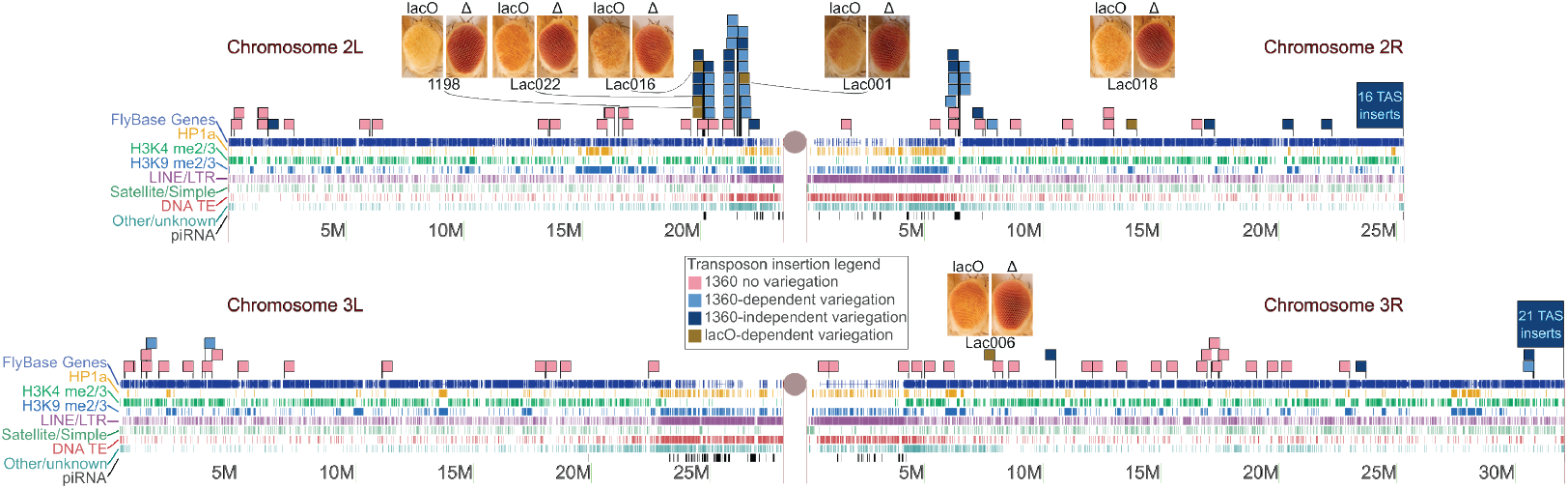
Identification of new insertion sites that support lacO repeat-dependent silencing. Scaled representations of chromosomes 2 and 3 (Release 6 plus ISO1 MT, NCBI RefSeq assembly GCF_000001215.4) with tracks illustrating the presence of annotated genes (FlyBase (Öztürk-Çolak et al. 2024)), HP1a binding (yellow), H3K4 methylation (green), and H3K9 methylation (blue) (modENCODE (Kharchenko et al. 2011)), algorithmically detected repetitive sequences, and piRNA clusters (Brennecke et al. 2007). Centromeres are represented by gray circles. Green flags illustrate the insertion sites of new lacO repeat lines described in this study. Other flags illustrate insertion of 1360-containing reporter elements from prior studies (Sentmanat and Elgin 2012; Huisinga et al. 2016) — red denotes full expression of the eye color reporter; purple denotes 1360 repeat-dependent variegation; blue denotes 1360 repeat-independent variegation. The latter includes high-frequency clusters of insertions in telomere-associated sequences (TAS); such insertions appear to be unique to 1360 and are not observed among lacO repeat transposon insertions. Pairs of photos for each location are shown with the left image representative of each line’s PEV phenotype and the right image representative of the phenotype after eye-specific expression of FLP recombinase to remove the lacO repeats.

### Mutations in H3K9 histone methyltransferases show little impact on *lacO* repeat-dependent silencing

The presence of HP1a and H3K9me2/3 and the absence of histone acetylation are prominent features of pericentric heterochromatin. In most animals HP1a and H3K9me2/3 are linked by a positive feedback loop, with methylated H3K9 recruiting HP1a, which acts as a histone reader and in turn recruits the histone writer SU(VAR)3-9, which di- and tri-methylates H3K9 (Bannister et al. 2001; Lachner et al. 2001; Schotta et al. 2002). The positive feedback of the system is believed to be important for heterochromatin spreading and maintenance during mitosis. *D. melanogaster* has three known H3K9 methyltransferases encoded by the genes *Su(var)3-9, eggless* (aka *egg* or *dSETDB1*) and *G9a*. These appear to operate in different chromatin domains; SU(VAR)3-9 plays a major role in pericentric heterochromatin, while *eggless* plays a major role in the heterochromatic fourth chromosome (Muller F element), and *G9a* on facultative heterochromatin and developmental chromatin remodeling (Mis et al. 2006; Stabell et al. 2006; Seum et al. 2007; Tzeng et al. 2007; Yoon et al. 2008; Brower-Toland et al. 2009; Merkling et al. 2015). While mutations in the genes for these three proteins act as dominant suppressors of PEV for reporters in their domain, they had little to no impact on *lacO* repeat-induced silencing (Figure 5A), suggesting that the ectopic heterochromatin induced by *lacO* differs from endogenous heterochromatin.

**Figure 5.**
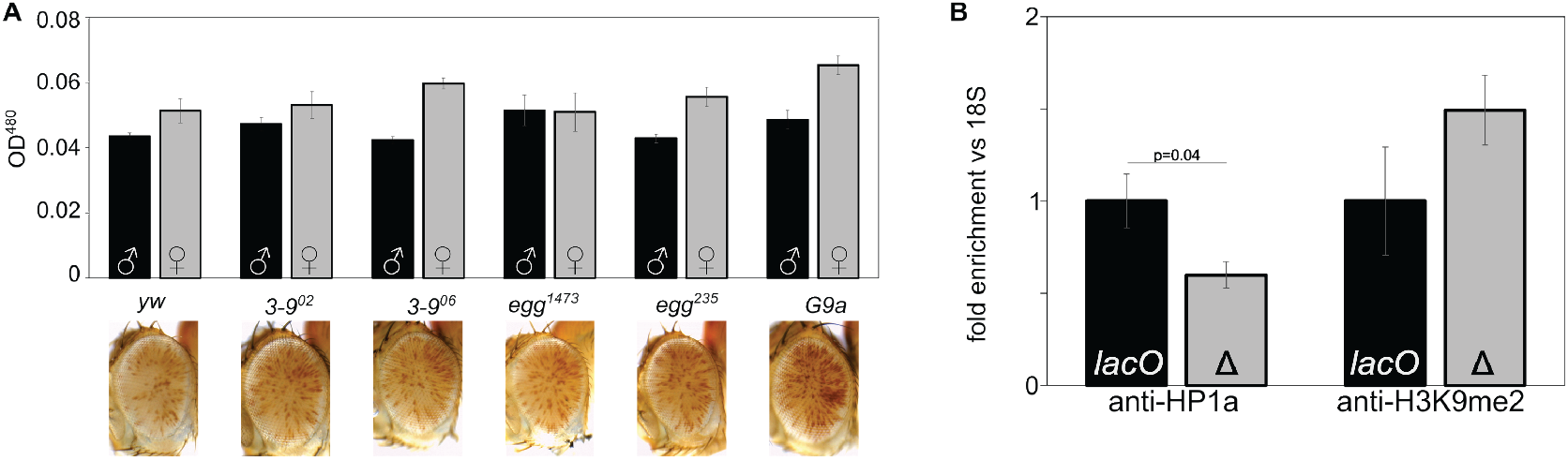
H3K9 di- and tri-methylation does not play a dominant role in line 1198-lacO repeat-dependent silencing. (A) 1198-lacO males were crossed to females carrying mutations in the three known H3K9 HMTs (Su(var)3-9, eggless, and G9a) and the progeny compared to controls (crossed to yw) with wild type HMTs (male progeny are pictured). Introducing these mutations had no effect on silencing as shown by examining the eyes (pictured) and determining pigmentation levels (black and gray bars for females and males, respectively). (B) Antibodies against HP1a (left) and H3K9me2 (right) were used for ChIP, followed by qPCR amplification of the 18S and hsp70 reporter promoters in mixed male and female adult flies with lacO repeats intact (lacO) or deleted (Δ). The HP1a and H3K9me2 levels are normalized to 18S relative to a lacO value of 1. Error bars denote standard deviation. HP1-binding is significantly reduced (p=0.04) after excision of lacO repeats while H3K9me2 is increased; the latter difference is of similar magnitude but is not statistically significant.

To further explore *lacO* repeat-induced heterochromatin we used ChIP-PCR to measure the amount of HP1a and H3K9me2 bound to the *hsp70* promoter of our reporter in line *1198-lacO* (see construct, Figure 1), comparing flies with the *lacO* repeats to those with the repeats removed by FLP-FRT recombination (Figure 5B). Deletion of the repeats reduces HP1a by about 40%, consistent with the loss of silencing observed in HP1a mutants. It is also associated with a 50% increase in H3K9me2, further supporting a decoupling of this mode of silencing and H3K9 di-methylation (Figure 5B). Thus, it appears that the silencing driven by the *lacO* repeats reflects an HP1a-dependent heterochromatin assembly that is uncoupled from the H3K9 HMT system.

### Histone de-acetylation plays a role in *lacO* repeat-induced silencing

To investigate the role of histone acetylation on *lacO* repeat-induced silencing, we tested the impact of nicotinamide, a small molecule histone deacetylase (HDAC)/sirtuin inhibitor with broad efficacy, feeding it to flies with one of three distinct PEV genotypes — our *1198*-*lacO* repeat line, the classic *white mottled 4* line (in which an inversion on the X chromosome moves the *white* gene adjacent to pericentric heterochromatin (Muller 1930; Solodovnikov and Lavrov 2022 July 1)), and a line in which an *hsp70-white* reporter is inserted into the pericentric heterochromatin of the fourth chromosome (Wallrath and Elgin 1995). Dietary nicotinamide suppressed variegation of all three PEV reporters (Figure 6A), indicating that, unlike the case of H3K9 methylation, *lacO* repeat-directed silencing shares a requirement for HDAC activity with other PEV reporters.

**Figure 6.**
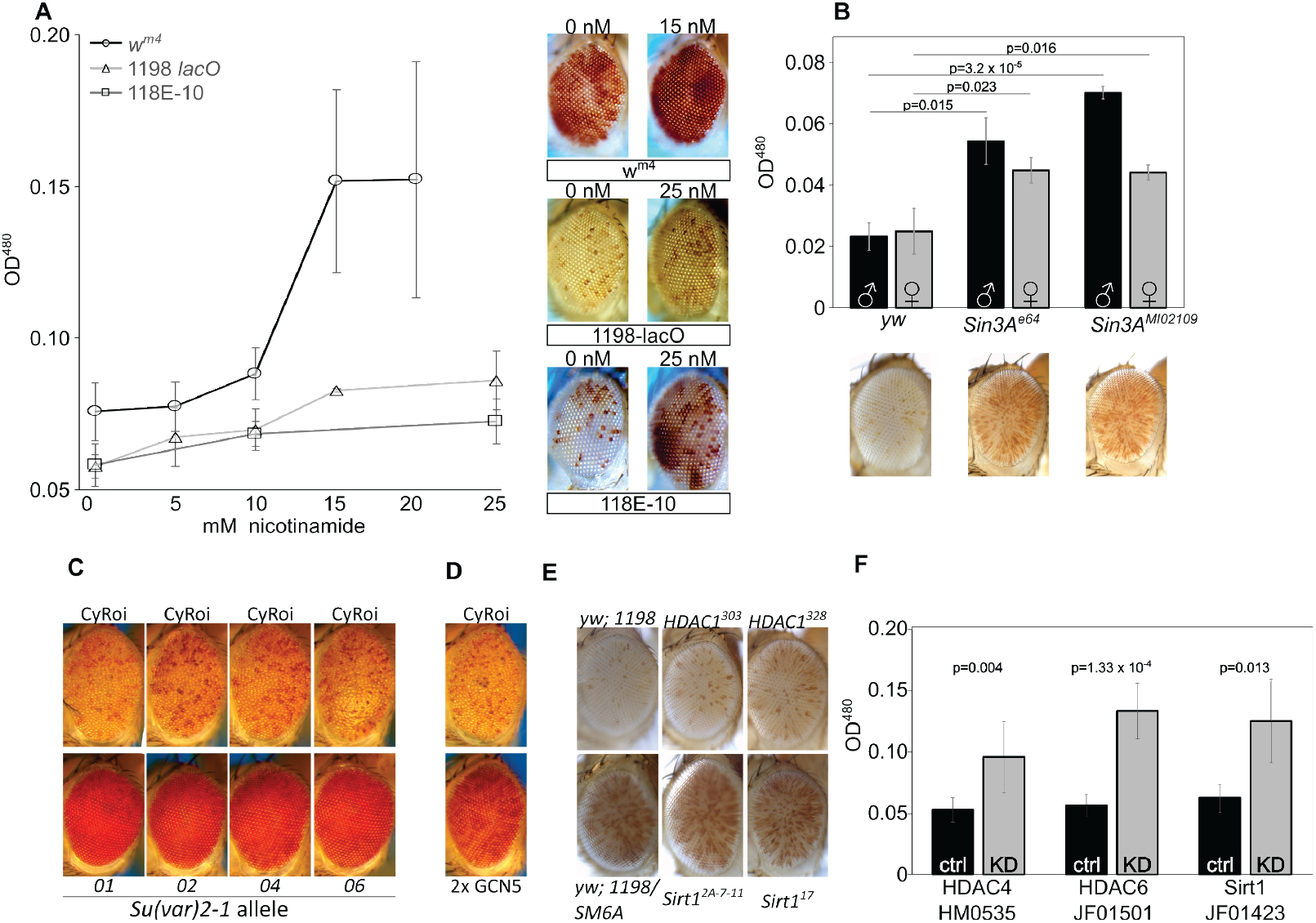
Histone deacetylation plays a major role in lacO repeat-induced silencing. (A) Mutant embryos from three different PEV lines were reared to adulthood on food containing the HDAC/Sir2 inhibitor nicotinamide, revealing a dose responsive suppression of variegation (males pictured; left 0 drug, right maximum drug tested). X-axis: millimolar concentration of nicotinamide; Y-axis : eye pigment, OD^480^. Line 118E-10 harbors a variegating reporter insert in Chromosome 4 heterochromatin, serving as a positive control for nicotinamide function. (B) Heterozygous Sin3A mutations suppressed variegation relative to wild type controls. 1198-lacO/CyO flies were crossed with yw (control) or with mutants for Sin3A and pigment analysis was carried out on adult progeny (females pictured; black bars = males, grey bars = females). (C, D) Heterozygous mutations in HDAC-interacting gene Su(var)2-1 (C) and overexpression of histone acetyltransferase GCN5 (D) both suppress variegation. E) Heterozygous HDAC1 and Sirt1 mutants have no significant effect on lacO repeat PEV. F) RNAi-mediated knockdown of HDAC4, HDAC6, and Sirt1 suppress variegation, further supporting a role for histone deacetylation in lacO repeat-directed silencing. Gray bars labeled “KD” express the driver and the indicated RNAi construct; black bars are no-RNAi controls.

We next investigated two proteins known to organize multi-subunit complexes that facilitate HDAC activity. Mutations in the HDAC-interacting gene *Sin3A* had a dominant suppressing effect (Figure 6B), further implicating histone deacetylation in *lacO* repeat-induced silencing. Similar results were seen with a series of mutations in the gene *Su(var)2-1* (Reuter et al. 1982) which facilitates histone demethylation (Yang et al. 2019) and deacetylation (Walther et al. 2020). Multiple EMS-induced *Su(var)2-1* mutants markedly suppressed *1198*-*lacO* variegation (Figure 6C). Further supporting the role of deacetylation, we observed suppression of silencing in flies with a transgenic extra copy of the *GCN5* lysine acetylase gene (Figure 6D).

We then sought to identify specific histone deacetylases that impact *lacO* silencing. Mutations in two well-studied histone deacetylases (Sirt1 and HDAC1) had no significant impact on variegation using the *1198*-*lacO* repeat line as reporter (Figure 6E). This result could be due to functional redundancy; there are five known histone deacetylases and five SIRT gene products with NAD-dependent histone deacetylase activity in *D. melanogaster* (Feller et al. 2015). Finally, we tested the effects of RNAi-mediated knockdown of histone deacetylases by crossing *1198*-*lacO* flies with UAS-driven shRNAs targeting HDAC and SIRT genes in the presence of a ubiquitous GAL4 driver (Figure 6F and Table S2). After an initial screen for suppression of variegation identified specific genes for further testing, follow up experiments with shRNAs targeting *HDAC4, HDAC6*, and *Sirt1* confirmed that knockdown of these gene products could suppress *lacO* repeat-induced variegation. It’s unclear why the heterozygous *Sirt1* mutation had no effect on variegation while *Sirt1* knockdown suppressed it; potentially the knockdown achieved lower protein levels. These results suggest that histone deacetylation is involved in *lacO* repeat-induced silencing.

We used ChIP followed by qPCR across the promoter region to measure H3K9 acetylation levels at our *1198*-*lacO* reporter gene and compared those levels to acetylation levels when the *lacO* repeats had been excised by FLP recombinase. Deletion of the *lacO* repeats leads to an increase in H3K9 acetylation (Figure 7) further confirming the importance of histone de-acetylation in establishment of the local *1198*-*lacO* repeat heterochromatin domain.

**Figure 7.**
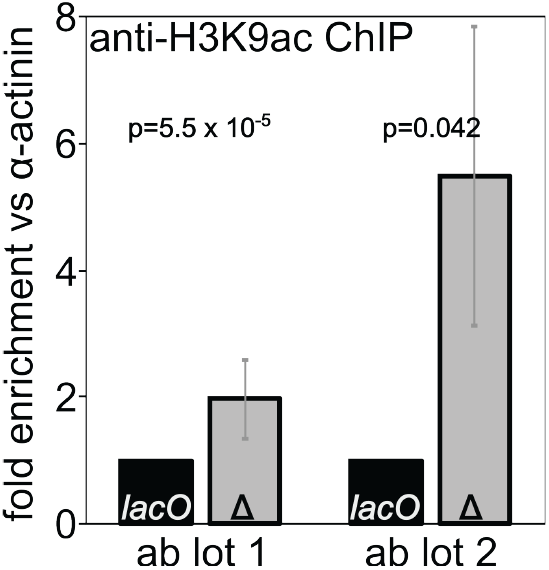
Deletion of lacO repeats increases H3K9 acetylation at the reporter gene promoter. Chromatin immunoprecipitation (ChIP) for acetylated H3K9 was performed on flies with (black bars) and without (gray bars) lacO repeats in the reporter. qPCR results for reporter promoter sequences were normalized to alpha-actinin promoter sequences, and the resulting values are expressed as fold difference between flies with the lacO repeats (lacO) and without the lacO repeats (Δ). Two lots of antibody gave qualitatively similar but quantitatively distinct results, in both cases supporting increased histone acetylation in the absence of the lacO repeats.

## Discussion

### A tandem array of 36 bp of *E. coli* DNA can drive HP1a-dependent silencing

Here we show that a 256 copy array of a 36 bp *E. coli lacO* DNA sequence inserted into the *D. melanogaster* genome can silence an adjacent reporter, apparently due to ectopic heterochromatin formation. Dependence on the tandem repeats is demonstrated by the complete loss of silencing upon their excision (Figures 1 and S4). In earlier screens using either *white* or *yellow* as a reporter within a P-element transposon devoid of repeats, variegation occurs only when the reporter is within or adjacent to a known heterochromatic domain such as the pericentric heterochromatin, the telomere-associated sequences (TAS), the F element (fourth chromosome), or the Y chromosome (Wallrath and Elgin 1995; Yan et al. 2002). In contrast, the variegating reporters we describe here are inserted into euchromatin. Thus we attribute their variegating phenotype to formation of an ectopic heterochromatin domain, driven by recognition of the tandem repeat of foreign DNA. This reveals a defense mechanism against intrusion by foreign DNA, as the *D. melanogaster* genome appears to have no prior exposure to this sequence, suggesting that the organism can defend itself against any tandem repetitious sequence (see Supplemental Discussion for additional references that informed our thinking).

The *lacO* repeat-induced variegation correlates with HP1a concentration, losing silencing in response to loss of HP1a function and gaining silencing when HP1a is overexpressed and tethered to the *lacO* repeats (Figures 2A and 2B). HP1a is associated with all constitutive heterochromatic domains identified in *Drosophila*, and is strongly correlated with reporter silencing. (Eissenberg et al. 1992; Eissenberg and Reuter 2009). Earlier studies from the Wallrath lab showed that tethering of HP1a to this *lacO* repeat array using an HP1a-lacI fusion protein is sufficient to induce silencing at a wide range of euchromatic insertion sites (Li et al. 2003; Danzer and Wallrath 2004). Association with HP1a may be sufficient to drive heterochromatin formation, either through interactions with other HP1 binding proteins such as SU(VAR)3-7 (Figure 2C) or through HP1a-driven phase separation (see Supplemental Discussion). *Our results here and several prior studies (e*.*g. (Honda et al. 2012)) imply a basic structural role for HP1a in heterochromatin assembly independent of H3K9 methylation* (Figure 5).

### An unusual temperature sensitivity for maintenance of the *lacO* repeat-induced silencing has allowed investigation of persistence of the licensing tag and PEV plasticity

In general, PEV is modestly suppressed at higher temperatures and enhanced at lower temperatures ((Gowen and Gay 1933; Hartmann-Goldstein 1967); reviewed in Spofford 1976; Girton and Johansen 2008)). In contrast, *lacO* repeat-induced variegation is robust at 25°C but fails at 18°C (Figures 3 and S4). This temperature-dependent plasticity operates in both directions (*i*.*e*., both gain and loss of silencing) throughout larval development up to pupariation (when the eye phenotype is fixed), indicating that at least one component of the *lacO* repeat-induced silencing system required for maintenance of heterochromatin fails at the lower temperature. This unexpected loss of silencing at 18°C allows an examination of plasticity in heterochromatin formation.

Generation of a stable heterochromatin domain in *Drosophila* is generally thought to occur in two major steps: initial uniform silencing in the early embryo followed by an ongoing “maintenance” stage. The initial silencing requires first a “recognition” step, identifying domains targeted for heterochromatin formation. The plasticity we observe, in conjunction with earlier observations, argues for a second “licensing” or marking step in late blastoderm/early gastrulation (nuclear cycle 14) that enables local heterochromatin formation, and maintains this memory through somatic cell division. Embryos must establish heterochromatin while activating the zygotic genome, and multiple lines of evidence identify the mid-blastula transition as the point at which pericentric heterochromatin is re-established via di- and tri-methylation of H3K9 and recruitment of HP1a (Yuan and O’Farrell 2016; Seller et al. 2019). Lu et al. proposed that the stochastic nature of PEV arises in the maintenance phase, with heterochromatin boundaries receding towards a point of origin prior to being fixed after terminal differentiation (Lu et al. 1996; Lu et al. 1998). In accord with this model, Gu and Elgin were able to show that Piwi, a component of the piRNA system used to recognize TEs for silencing (reviewed in (Czech et al. 2018)), is required in the early embryo but not in the larval stages for silencing of a PEV reporter (Gu and Elgin 2013), indicating that components of the initial recognition step are not needed after licensing. In contrast, they found that HP1a is required both in the early embryo, potentially as part of the licensing step, and subsequently to maintain silencing. Similar results pointing to a key role for HP1a in the initial establishment of heterochromatin in the early embryo (licensing) have also been reported (Bughio and Maggert 2022).

Two complementary lines of evidence narrow the temperature-sensitive component of *lacO* repeat-induced silencing to the maintenance phase. When larvae are moved from 25°C to 18°C, loss of silencing is observed implying a failure of heterochromatin maintenance (Figure 3A). Conversely, when larvae are moved from 18°C to 25°C, restoration of silencing is observed (Figure 3B), implying that the recognition and licensing steps are both functioning properly to mark *lacO* repeats for silencing even when the adult phenotype is suppressed. In other words, the license persists even when the silencing does not. In contrast, the Gu and Elgin (2013) study showed that the depletion of HP1a in the early embryo resulted in loss of silencing in the adult eye, even when normal levels of HP1a were restored during zygotic gene expression. *Combining these observations leads to the conclusion that HP1a plays a key role in licensing, which allows the assembly of heterochromatin at the marked site when the shift to 25°C occurs during the larval stage, permitting heterochromatin assembly*.

### A small subset of reporter insertion sites supports *lacO* repeat-dependent ectopic heterochromatin formation

The studies reported here primarily used a single insertion site previously shown to exhibit TE-dependent silencing (Sentmanat and Elgin 2012). Mobilizing the P element *lacO* reporter construct revealed additional repeat-dependent silencing sites, 5 of which we mapped; all of these displayed the same temperature- and repeat-sensitivity as the original *1198*-*lacO* line (Figures 4 and S4, and Table 2). Several studies using the transposon remnant *1360* (Sentmanat and Elgin 2012; Huisinga et al. 2016) or the *lacO* array (Li et al. 2003; Danzer and Wallrath 2004) have found that insertion of the repeats themselves is not sufficient to induce ectopic heterochromatin formation at most sites; the vast majority of the inserted repeat-containing P elements observed in transposition screens, including this one, are distributed throughout the genome and exhibit wild type red eye color. Here we uncover sites that are normally euchromatic (permissive for reporter expression) but can be converted to heterochromatin by the insertion of the *lacO* repeat array. The small proportion of insertion sites resulting in variegating eye color are strikingly clustered, indicating that this phenomenon is restricted to a small number of chromosomal subdomains, illustrated by the clustering of insertion sites that meet these criteria (Figure 4). Combining multiple studies of *lacO* repeat- or *1360-*dependent PEV lines from our labs, twenty-one out of 33 inserts (64%) where variegation is confirmed to be repeat-dependent fall into two clusters within a single 1.6 Mb region of chromosome arm 2L that constitutes only 1.3% of the 120 Mb euchromatic genome. Another six (18%) can be found in a 300 kb segment of arm 2R, 0.25% of the euchromatic genome. Thus >80% of all confirmed cases of repeat-dependent variegation using either TE remnant *1360* or *lacO* inserts fall within the same 1.5% of the euchromatic genome. The two genomic regions identified are characterized by their proximity to repetitive DNA and heterochromatin blocks, but not uniquely so. *Other factors which we have not been able to identify must be contributing to the formation of stable heterochromatin in response to the presence of multiple distinct types of repetitive DNA. This indicates that the feature(s) that define these domains are linked to licensing and maintenance, not recognition*.

How might a tandem array of a novel foreign DNA sequence be recognized as a target for silencing? Heterochromatin tends to be AT-rich, and experiments inserting bacterial DNA into yeast chromosomes have suggested that differences in AT content might play a determining role (Meneu et al. 2025). In the case of the endogenous tandem repeats in *D. melanogaster*, multiple mechanisms appear to be in play. Examples of repeats in *D. melanogaster* range from minisatellites (tandem arrays with a repeat of a few nucleotides) to the histone gene cluster (~110 copies (Crain et al. 2024; Shukla et al. 2024)) and the rRNA genes (80-600 copies (Guetg et al. 2010; Lu et al. 2018)), the latter two both repeating on a kilobase scale (Watase et al. 2022; Ahmad et al. 2025). Several studies have identified specific proteins that bind to these endogenous repetitious sequences and appear to be critical for driving heterochromatin assembly: for example D1 in the case of AT-rich satellite DNAs (Aulner et al. 2002; Blattes et al. 2006), HERS in the case of the histone genes (Ito et al. 2012), and NoRC in the case of rRNA genes (Guetg et al. 2010). However, these strategies do not appear to be well-suited for dealing with novel invading DNA repeats. The *lacO* fragment has a similar GC% as the rest of the *D. melanogaster* genome, no significant similarity to any endogenous sequences or piRNAs, and no regions of mono-di- or tri-nucleotide repeats that might engage known silencing mechanisms. There is no reason to anticipate the presence of a eukaryotic nuclear protein that can specifically recognize the *lacO* repeat as a foreign invader to be silenced.

Evidence from multiple systems implicates pausing by RNA polymerase (e.g., *Homo sapiens* (Colak et al. 2014; Groh et al. 2014; Loomis et al. 2014); *Caenorhabditis elegans*, (Zeller et al. 2016)) or DNA polymerase (Gaggioli et al. 2023), leading to formation of R-loops and/or D-loops at repetitive sites as a trigger for epigenetic silencing. While some of these mechanisms appear to rely on H3K9 methylation, loop formation can also cause DNA damage and recruitment of DNA repair and heterochromatin proteins, notably HP1a and HDAC1/2 (e.g., (Luijsterburg et al. 2009; Miller et al. 2010; Zeller and Gasser 2017), and reviewed in (Lemaître and Soutoglou 2014)). Such a mechanism is an attractive candidate to explain *lacO* repeat-induced silencing because it does not depend on a specific DNA sequence, relying instead on the inherent propensity of repetitive transcripts to form RNA:DNA hybrids (reviewed by (Zeller and Gasser 2017)).

*We conclude that the lacO-repeat array will drive silencing by heterochromatin formation, resulting in a variegating phenotype in an adjacent reporter, in a small number of chromosomal domains. One possible mechanism for recognition of the repeat array is the formation of R-loops and/or D-loops. If this is the case, the lacO-repeat insertion sites that trigger heterochromatin packaging must allow accessibility for transcription but also encompass unidentified features that predispose to repeat-dependent silencing*.

### Heterochromatin induced by the *lacO r*epeat has a distinct biochemistry

The heterochromatin structure detected here, tracked primarily by its ability to induce silencing of a reporter gene, is dependent on the core heterochromatin proteins HP1a and SU(VAR)3-7 (Figure 1) and on histone deacetylation (Figure 6), but not on H3K9 methylation (Figure 5). This is surprising given that H3K9 di- and tri-methylation is a prominent feature of pericentric heterochromatin as well as silenced TEs (Penke et al. 2016), trinucleotide repeats (Nageshwaran and Festenstein 2015), and tandem gene arrays (van Steensel et al. 2001). A study that parallels our experiments with the *lacO* repeats in flies reported here, starting from the 1198 construct but inserting the GAA repeat from a Friedrich’s Ataxia patient, also shows a requirement for H3K9 methylation for silencing (Gracheva et al., manuscript in preparation).

While a heterochromatic silencing system based on localization of HP1a complexes and histone deacetylation is not the predominant model, it is not without precedent. Similar examples have been reported in other systems. For example, HP1-induced ectopic heterochromatin that is H3K9-independent and HDAC-dependent has been reported *in Neurospora crassa* (Honda et al. 2012; Gessaman and Selker 2017). A recent study in *C. elegans* showed that the HP1a homologue HPL-2 functions with LIN-13 (a nuclear regulatory protein) to promote heterochromatic foci and gene repression independently of H3K9 methylation (Delaney et al. 2025). There is also evidence for the parallel presence of an HMT-independent mechanism for HP1a recruitment at the 359 bp satellite sequence in *D. melanogaster* (Yuan and O’Farrell 2016); this system is consistent with the observations here, suggesting a heterochromatin based on the presence of HP1a and maintenance of de-acetylated histones (Figures 2A, 2B, and 6).

Many studies have identified histone deacetylation as a key early step in the transition from a euchromatic to a heterochromatic state: in cases where H3K9 methylation is a requirement, H3K9 deacetylation appears to be a prerequisite (reviewed in (Elgin and Reuter 2013)). Indeed, broad deacetylation has been suggested to be a necessary step upstream of heterochromatin formation in the *Drosophila* embryo (Walther et al. 2020) and in *S. pombe* (Alper et al. 2013). Although analyzing histone acetylation can be difficult in flies due to redundancy among the five known HDACs (Pallos et al. 2008) and five known Sirts, we were able to generate several lines of evidence indicating that histone deacetylation is a key step in generating the local heterochromatin found at the *lacO* repeat site (Figure 6). The *lacO* repeat-induced silencing was reversed by nicotinamide, a broad suppressor of HDAC/sirtuin activity (Figure 6A); by mutations in *Sin3A* or *Su(var)2-1*, both organizers of HDAC complexes (Figures 6B and 6C); by overexpression of GCN5, a histone acetyltransferase (Figure 6D); and by knock-down of specific HDACs (Figure 6E). Su(var)2-1, initially identified as a suppressor of variegation (Reuter et al. 1982), was recently shown to recruit both HDAC1 (Walther et al. 2020) and the H3K4me1/2 demethylase SU(VAR)3-3 (Yang et al. 2019) to sites of transcriptional repression. Sin3A is also part of a protein complex with HDAC1, and interacts physically and genetically with H3K4me3 demethylase *KDM5* (Spain et al. 2010; Gajan et al. 2016) to promote silencing. Together these results argue for a primary role for histone deacetylation in promoting silencing.

In sum, this study of the impact of foreign tandem repeats on local gene expression has identified a form of heterochromatin with distinct features, including distribution in the genome, temperature sensitivity, and biochemistry. These features can now be exploited to further our understanding of the formation, specificity, and stability of heterochromatin and its role in genome integrity.

## Materials and Methods

### Fly husbandry

All flies unless otherwise indicated were cultured at 25°C in 60-70% relative humidity. Most experiments used a fly food recipe consisting of 45 g agar, 155 g brewer’s yeast, 674 g sucrose, 858 g cornmeal, 8.5 L water, 100 mL propionic acid, and 70 mL of 10% tegosept. Generation and characterization of the *lacO* transposition mutants was carried out on the standard sucrose cornmeal media recipe used by the Bloomington *Drosophila* Stock Center with a sprinkle of dry yeast pellets.

### Generation of the *1198-lacO-hsp70-white* stock

Construction of the original *1360* hsp70 w+ P element reporter P{T1} has been described previously (Haynes et al. 2006). To swap *1360* for the *lacO* array the *loxP-frt* elements from vector *pCR2*.*1-loxP-frt* were cloned into the *XhoI* site of *pCR2*.*1-attB1-loxP-y-attB2* using primers *XhoI8-24F/R* to make *pCR2*.*1-attB1-loxP-y-loxP-frt-attB2* (Sentmanat and Elgin 2012). The *lacO* cassette is comprised of 256 copies of a 36 nt *E. coli* lac operator sequence CCACATGTGGAATTGTGAGCGGATAACAATTTGTGG (Sasmor and Betz 1990) and its cloning has been previously described (Li et al. 2003). The repeat array was excised from pBS KS(-)-*lacO* using *XhoI* and inserted into *pCR2*.*1-attB1-loxP-y-loxP-frt-attB2* to make *pCR2*.*1-attB1-loxP-y-loxP-frt-lacO-attB2*. The size of the insertion was confirmed by enzymatic digest and agarose electrophoresis. Male 1198-*1360* flies (Sentmanat and Elgin 2012) were crossed to females expressing the phiC31 integrase in germline cells driven by the *vasa* promoter (BDSC 24483) and embryos were injected with the *lacO*-containing vector. Putative F1 male recombinants were selected by the presence of the *yellow*+ phenotype and full cassette exchange was verified by PCR (Figure S1) as previously described (Sentmanat and Elgin 2012).

### Homology search for *lacO* sequences in *D. melanogaster* genome

NCBI blastn (Altschul et al. 1997) was used to compare the 36 nucleotide *lacO* sequence (CCACATGTGGAATTGTGAGCGGATAACAATTTGTGG) and a 256-copy array of that sequence to the Release 6 plus ISO1 MT assembly (RefSeq accession GCF_000001215.4) maintained by the FlyBase consortium (Öztürk-Çolak et al. 2024)), and to a recent PacBio HiFi assembly (GenBank accession GCA_042606445.1) (Shukla et al. 2024). Parameters were: Expect threshold = 0.05, Word Size = 11, Match/Mismatch Scores: 2,-3, Gap Costs: Existence: 5 Extension: 2, Low complexity regions filter = off.

### *In vivo* excision of *lacO* repeat array

The *lacO* repeats are flanked by FRT sites (Figure 1), allowing precise removal of the repeat array from the rest of the reporter construct in live flies expressing the FLP recombinase. Creation of stable repeat-deletion lines has been described previously (Sentmanat and Elgin 2012). Briefly, *1198-lacO* males were crossed to females expressing a heat shock-inducible FLP on the X chromosome (BDSC 8862), and progeny were incubated at 37°C for 1 hour on days 3 through 7 of larval development. F1 males with mosaic eye phenotypes suggesting high levels of FLP-mediated recombination were crossed to second chromosome balancer stocks and individual F2 males were selected to create stable balanced stocks — excision of *lacO* repeats was verified by PCR using primers A412F and 200R (Haynes et al. 2006).

### Testing of dominant modifiers of *lacO*-directed silencing

Candidate variegation-modifying mutant flies were crossed with *1198-lacO/CyO* flies in plastic bottles, approximately 20-25 females and 10-15 males. The presence of mutants was deduced by the absence of dominant balancer phenotypes and dominant eye effects were assessed qualitatively by visual comparison of pigment levels between mutants and balancer, and between mutant and a parallel cross to *yw*^*67c23*^ which does not carry a known PEV modifier and is treated as wild type for the purposes of these studies. For most experiments we crossed mutant males (for example *Spn-E, mael, AGO2, E(z)*, or *HP2*) with *1198-lacO/CyO* females. To account for the possibility of maternal effects in chromatin mutant lines (Gu and Elgin 2013) we crossed mutant females with *lacO* males.

### RNAi Knockdown Cross Protocol

RNAi-driven knockdown of a variety of genes of interest was done using stocks from the Transgenic RNAi Project (TRiP). 10 female flies with 1198-*lacO* and a ubiquitously expressed GAL4 driven by the *daughterless* promoter (*yw;* 1198*-lacO/CyO; da-GAL4/Tm3 Sb*^*1*^) were crossed to 5 balanced or homozygous males with GAL4-driven RNAi hairpins targeting specific genes of interest. As a control, 2 TRiP lines (*HDAC4*^*HM05035*^ and *Su(z)12*^*HMS00280*^) were crossed to 1198*-lacO* flies without the *da-GAL4* driver. To eliminate confounding X chromosome markers common to all TRiP stocks only male progeny with no balancer alleles were selected for phenotypic analysis. In a subsequent version of the experiment, each TRiP line was crossed to *lacO* flies with and without the da-GAL4 driver to provide more consistent controls.

### Fly Eye Imaging

Two distinct eye imaging protocols were used in this study. For all except Figure 4, *Drosophila* adults were randomly collected within 24h after eclosion and aged for 3-4 days at 25°C for full pigment development. They were anaesthetized and mounted on microscope slides. Pictures of eyes were taken with a digital camera installed on a Leica S8APO dissecting microscope and processed using ImageJ software. Initial pictures were taken using a white background. Later pictures, especially for flies utilizing RNAi knockdown, used a blue background for greater contrast with the red eye pigment. Detailed methods for Figure 4 are described in (Pipkin et al. 2024). Briefly, 3-4 day old adult progeny were frozen overnight and photographed using a full frame digital camera and macro lens mounted on a horizontal focus stacking rail. Images were acquired in RAW format with exposure of 1/25, f2.8, ISO 500. Rail travel from top to bottom is 2750 uM made up of 56 steps at 50 uM each. The 56-image stack was automatically exported from Helicon Remote to Helicon Focus version 7.7.5 and a composite image combining the most-focused pixels of each individual image was generated using the “C, smoothing 4” setting and saved as TIF files.

### Eye Pigment Assay and Statistical Analysis

In *white* mutant flies, eye pigmentation reflects expression of the transgenic white reporter gene. Spectrophotometric measurement of eye pigments (described in (Sun et al. 2004)) were made on pools of 15-25 3-day old adult flies from at least 4 crosses, and carried out as described in (Sun et al., 2004) with minor changes. Flies were mechanically homogenized in Pigment Assay Buffer (0.01 N HCl in ethanol), followed by incubation at 50°C for 10 minutes, centrifugation, and measurement of 480 nm absorbance of the supernatant homogenate to quantitatively determine the level of red drosopterin pigments. We applied a homoscedastic Student’s t-Test in Microsoft Excel to generate 2-tailed p-values and, where appropriate, adjusted for multiple comparisons by multiplying the p-value by the number of tests within a given panel. All bars show the average of at least three biological replicates and error bars show standard deviations.

### ChIP Protocol

Chromatin immunoprecipitation followed the modENCODE protocols (Kharchenko et al. 2011; Riddle et al. 2011) from 300-1000 mg of whole 4-day old adult 1198 *lacO*/CyO or 1198 Δ/CyO flies. Each ChIP experiment included at least two independent experimental replicates, and at least two technical replicates for each sample. Chromatin was sheared over six rounds of 30 sec on/ 30 sec off in Bioruptor (Diagenode) resulting in average fragment sizes of ~100-200 bp and immunoprecipitated with antibodies to HP1a (W191) and H3K9me2 (Abcam 1220). The relative enrichment of each mark at the designated region was determined by quantitative PCR (iQ SYBR Green Supermix, Bio-Rad) for 18S and *hsp70-white*, using 18S to normalize *hsp70-white* between the two genotypes. Two replicates of each PCR were run and input control values were subtracted before calculation of the cycle threshold from the average of the replicates.

The transient nature of histone acetylation required modification to standard protocols. To preserve chromatin acetylation state, the PBS-EDTA+, nuclear extraction buffer, and wash buffers were supplemented with 0.01 mM TSA and 10 mM nicotinamide. Immunoprecipitations were performed using a mouse monoclonal antibody against H3K9ac (MAB Institute, 309-32379, Lots 13012 and 16009). Mock IPs were performed using mouse IgG (Jackson ImmunoResearch Laboratories, 015-000-003, Lot 150101). DNA extracted from immunoprecipitated chromatin fragments was subjected to three real-time PCR analyses using primers for *hsp70-white* (above) and for *α-actinin*. Each PCR experiment included H3K9ac IP, mock IP, and input DNA from *1198 lacO/CyO* or *1198 Δ/CyO* flies. Acetylation enrichment levels were determined based on input-normalized PCR results obtained for the *hsp70-white* region, and then normalized again to mock IP. Acetylation values obtained for 18S was used to normalize *1198-lacO* and *1198 Δ* for comparison.

### The *lacO* transposition mutagenesis screen

To facilitate a transposition mutagenesis screen for genomic locations on the autosomes that support *lacO* repeat-mediated silencing we first needed to mobilize the 1198 *lacO* P element and recover insertions on the X chromosome. Unmated *w*; *sp*/CyO; *delta2-3 sb*/TM6B (BDSC 3612) females which express the P element transposase were crossed with *w*; 1198 *lacO/CyO* males. Female F1 *lacO/CyO*; *delta2-3 sb* were crossed to *w*; *net*; *sbd*; *spa* males homozygous for viable recessive markers on all autosomes. Individual male F2 progeny with CyO (indicating the absence of the parental *lacO* insert) and eye color (indicating the presence of a novel *lacO* insertion) and without *stubble* (indicating absence of the transposase) were back-crossed to multiply marked recessive females. Stocks with red eyes and all three recessive phenotypes were presumed to carry a new *lacO* insertion on the X chromosome. Two X chromosome *lacO* stocks were recovered, and the presence of the full length *lacO* insert was confirmed by Southern blot (see Supplemental Data) and mapped by inverse PCR as previously described (Huisinga et al. 2016). Southern blots were performed using 10 µg of total genomic DNA from the stocks of interest, hybridized to probes labeled with DIG-High Prime DNA Labeling and Detection Starter Kit II (Millipore Sigma/ Roche 11585614910) according to the manufacturer’s instructions. Inverse PCR mapping was performed as described in (Sun et al. 2004) and Southern blots as described herein.

Insertion of the full length *lacO* construct was confirmed by Southern blot (Figure S2). Total DNA was extracted from about 100 adult flies in 500 µl of lysis buffer (0.1 M EDTA, 0.1 M Tris, 1% SDS, 1% DEPC), incubated 30 min at 75°C. 70 µl of 8 M potassium acetate was added to homogenate, mixed vigorously, and incubated on ice for 1 h before 10-minute spin at top speed in a desktop centrifuge, phenol/chloroform extraction, isopropanol precipitation, and 70% ethanol wash. Approximately 10 µg of genomic DNA was incubated with HindIII, EcoRI, and XhoI restriction endonucleases and separated by electrophoresis on 0.8% agarose. DNA transfer to PVDF membrane and membrane hybridization was performed as described in manufacturer’s instructions for DIG-High Prime DNA Labeling and Detection Starter Kit II (Roche 11585614910). Hybridization probe for the transgenic *yellow* sequence in our *lacO* cassette was produced by PCR using primers Pvu yellow F and Bgl yellow R. These newly isolated X chromosome *lacO* lines were used for subsequent transposition screens to recover lines exhibiting *lacO*-repeat induced silencing; the insertion sites were mapped by inverse PCR as previously described (Sentmanat and Elgin 2012).

We used a custom mirror of the UCSC genome browser (https://gander.wustl.edu/) to assess the proximity of the insertion sites to transposons in the *D. melanogaster* “Aug. 2014 (BDGP Release 6 + ISO1 MT/dm6)” genome assembly (NCBI accession GCF_000001215.4 (Hoskins et al. 2015)) and to create map displays of insertion sites relative to other genomic features. Genomic regions surrounding the insertion sites were shown with the “Combined Repeats” evidence track showing on the which shows the regions of the *D. melanogaster* genome with similarity to sequences in the “centroid” *Drosophila* repeat library that have been detected by RepeatMasker (Smit et al. 2013) using the following parameters: -nolow -s -e wublast. The “centroid” *Drosophila* repeat library consists of *Drosophila* transposons in release 20150807 of the RepBase repeat library (Jurka et al. 2005), *Drosophila* helentrons (Thomas et al. 2014), and *Drosophila ananassae* transposons identified by five *de novo* repeat finders. The five de novo *D. ananassae* repeat library include sequences from the ReAS repeat library produced by the *Drosophila* 12 Genomes Consortium (Clark et al. 2007), LTRHarvest (Ellinghaus et al. 2008), RepeatModeler (Smit and Hubley 2008), Tedna (Zytnicki et al. 2014), and dnaPipeTE (Goubert et al. 2015). The protocol used to construct the “centroid” *Drosophila* repeat library has previously been described ((Leung et al. 2017) in Supplementary File S7 pages 22–25). To create the maps in Figure 4 we used the same UCSC browser instance displaying tracks for FlyBase genes, modENCODE ChIP Data for Histone Modifications (H3K4 and H3K9 methylation), modENCODE ChIP Data for Chromosomal Proteins (HP1a/Su(var)205), piRNA clusters (Brennecke et al. 2007), and the repeat detection tracks described above.

As described above, the *lacO* repeats are flanked by FRT sites to allow FLP recombinase-mediated excision of the repeat cassette. Repeat-dependence of variegation in newly isolated *lacO* lines was tested by crossing them to flies expressing eye-specific FLP recombinase driven by the *eyeless* promoter (*yw; eyFLP/TM3, yw; eyFLP/Sb*, or *yw; eyFLP Sb/TM3*, all derived from BDSC 5576) – silencing was considered repeat-dependent if *eyFLP* caused a substantial suppression of variegation relative to siblings without *eyFLP*. To assess temperature dependence of repeat-induced silencing, newly isolated *lacO* lines were crossed to *yw* at 18°C or 25°C for 24 hours and 2-4 day old adults were collected and photographed, and eye pigment levels were measured and compared as described in (Pipkin et al. 2024). Data from these experiments are shown in Figure S4.

### Temperature shift experiments

Stocks were propagated at the experimental starting temperature in standard media for several generations prior to starting the temperature shift experiments. Roughly 10 female and 3 adult male *yw; 1198-lacO/CyO* flies were placed in vials were placed in vials of fresh food for 48h and transferred for 24 hours to an enriched semi-defined media containing 10 g agar, 80 g brewer’s yeast, 20 g yeast extract, 20 g peptone, 30 g sucrose, 60 g glucose, 0.5 g MgSO_4_ x 6H_2_O, 0.5 g CaCl_2_ x 2H_2_O, 6 mL propionic acid, 10 mL 10% p-Hydroxy-benzoic acid methyl ester in 95% ethanol in 1 L distilled water (BDSC Semi-Defined Food). For each timepoint 3 independent vials were moved from the starting temperature to the shifted temperature for the remainder of development. Timepoints were 24 hours (embryo group), after hatching of first instar larvae (L1 group), after most larvae had entered the wandering stage (L3 group), and after most larvae had formed pupae (Pupal group). The control group remained at the starting temperature. For experiments starting at 18°C we increased the number female parents to compensate for decreased rate of egg laying at the lower temperature.

### Nicotinamide feeding experiments

Nicotinamide is water soluble and was added as powder to warm (~50°C) semi-defined fly food to make a concentrated stock of 100 mM. This 100 mM nicotinamide food was diluted with unmodified food to final concentrations of 10 mM, 15 mM, 25 mM. 5 female and 2-3 male flies from each PEV stock was transferred to fresh medium for 48 h before transfer to enriched semi-defined medium containing nicotinamide. Parents were discarded after 24 h and F1 adults were collected, aged, sexed, and subjected to pigment assays upon eclosion.

### Isolation and testing of novel Su(var)2-1 and Su(var)3-7 alleles

To isolate novel Su(var) mutations we carried out a mutagenesis screen in *w*^*1118*^ males (BDSC 3605) on 2.5 mM ethyl methanesulfonate (EMS) crossed to PEV flies with a sensitized genetic background containing a spontaneous enhancer of variegation mutation (E(var)1^01^) on a multiply marked balancer. Mutagenized males were crossed with *In(1)w*^*m4*^; *T(2;3)ap*^*Xa*^ *+ In(2L)Cy, ap*^*Xa*^ *Cy E(var)3-1*^*01*^. The presence of the enhancer of variegation allele E(var)3-1^01^ silences the *w*^*m4*^ allele leading to absence of eye color – even modest suppression of variegation can be detected on this background (Reuter and Wolff 1981; Reuter et al. 1985; Wustmann et al. 1989), and flies with this phenotype were isolated and sequenced as described in (Walther et al. 2020). The impact of these novel alleles on *1198-lacO* were tested by crossing *1198-lacO* females with *w*^*m4*^; *Su(var)2-1*/CyRoi or *w*^*m4*^; *Su(var)3-7*/TM3 Sb Ser males and comparing eye color between mutant and balancer male siblings.

### GCN5 overexpression

To test the impact of overexpression of GCN5 on *lacO*-mediated silencing we injected the *P*{*FlyFos028109-Gcn5-V5-3xFLAG*} vector containing a transgenic copy of *Gcn5* under the control of the endogenous *Gcn5* promoter into embryos with a 2nd chromosome attP docking site for phiC31 integrase-mediated transformation (BDSC 9722). Transgenic flies were generated according to (Bischof et al. 2007; Ejsmont et al. 2009) and males were crossed to *1198-lacO* females.

## Supporting information

Supplemental Data and Discussion

## Acknowledgements

We thank Michael Grupe (Washington University in St Louis) for checking the integrity of the P-element construct after transposition (Figure S2) and Emily Chi (Washington University in St Louis) for initial examination of the temperature effect. We are grateful for the essential work screening and characterizing novel *lacO* mutants by undergraduate students in Gene Expression, Molecular Techniques, and Advanced Research Projects classes at Bemidji State University. We thank Professor Thomas Jenuwein for support of the *Drosophila Su(var)* project and Maria Kube and Ramona Abe for experimental support. We thank members of the Elgin, Reuter, and Arsham research groups for discussion, feedback and general support. Thanks to Lori Wallrath for providing the *lacO* repeat fragment. We’re grateful to members of the generous and vital *Drosophila* research community for supplies and discussion.

