## Supplemental Data and Discussion for "Heterochromatin-based silencing of a foreign tandem repeat in *Drosophila melanogaster* shows unusual biochemistry and temperature sensitivity"

### PCR Primers

Table S1

| | |
|---|---|
| *lacO* cloning | |
| XhoI8-24 F | CTGGCGGCCGCTCGACAGGAAACAGCTATGACCATGA |
| XhoI8-24 R | TAGATGCATGCTCGACTATAGGGCGAATTGGGCCCTC |
| cassette exchange confirmation | |
| attB2 F | GTGGATCCACTAGTTCTAGAGC |
| white R | ATTGATGGCGTAAACCGCTTGGAG |
| 1198 F | GGCATTGAATGAGCATTGTAATCGATACTT |
| attB4/09 R | ATCAAGCTTATCGATACCGTCGACC |
| Y 5’ R | TCAAGCGACCAGGCGATCTCAAAT |
| ChIP | |
| 18S F | TTCATGCTTGGGATTGTGAA |
| 18S R | GTACAAAGGGCAGGGACGTA |
| hsp70 F | CAAGCGCAGCTGAACAAGCTAAAC |
| white R | ATTGATGGCGTAAACCGCTTGGAG |
| *α-actinin F* | CAGCAAGCACCTCTGCTCTA |
| *α-actinin R* | TGCAAGCGTTAGTGAGATCC |
| yellow hybridization probe for southern | |
| Pvu y F | TATCGTCCTGTCTTGCCACA |
| Bgl y R | ACTTTCCCTGCACCCAAA |
| *lacO* repeat excision confirmation | |
| A412F | CGTTAACGTTAACGTTCGAGG |
| 200R | GACCTACCACAATAACCAGT |

### PCR Verification of Recombination-Mediated Cassette Exchange replacing *1360* with *lacO*

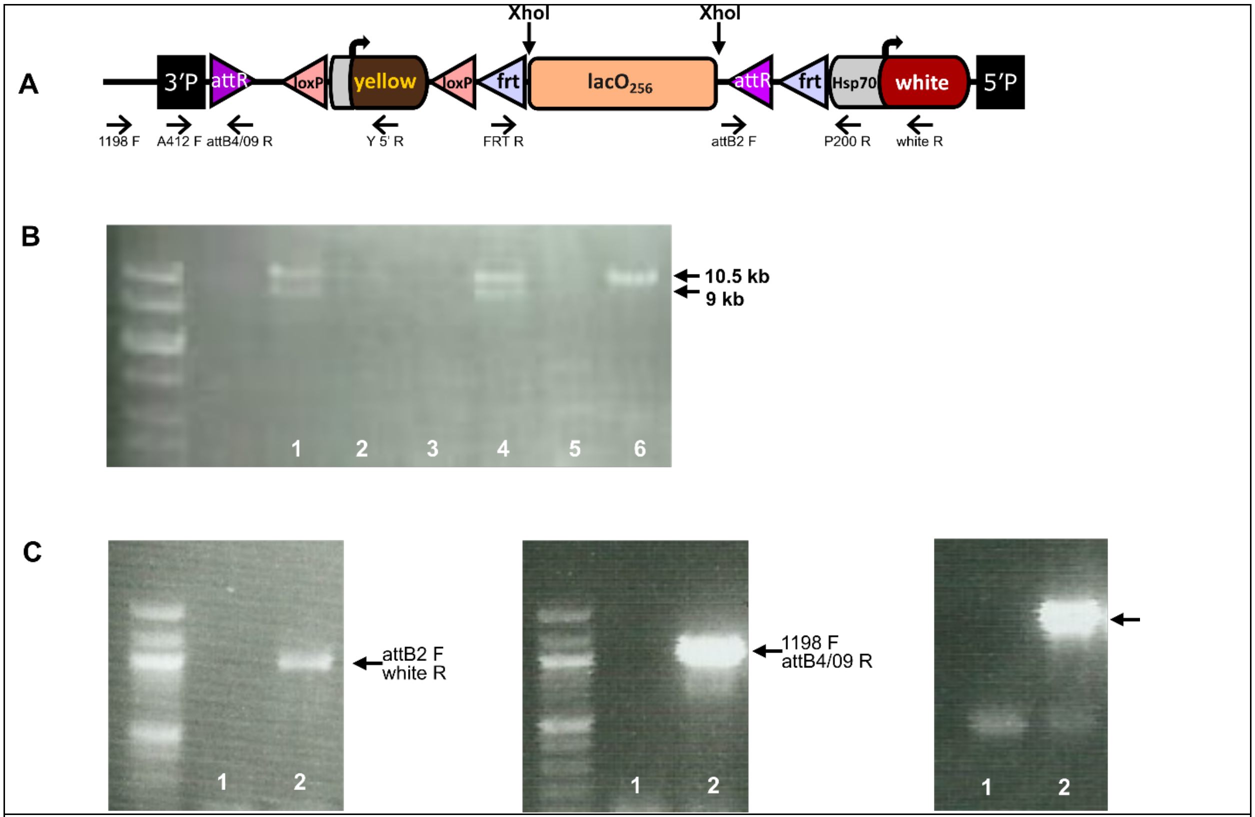


*Figure S1. Confirmation of the* lacO *insertion in the RMCE genomic landing. A. Illustration of the final* 1198-lacO *P element reporter after genomic integration and cassette exchange, including 3' flanking genomic sequence. The reporter was derived from the pCR2.1-attB1-loxP-y-loxP-frt-attB2 vector. The* lacO *fragment is flanked by two XhoI sites. B. pCR2.1-attB1-loxP-y-loxP-frt-lacO-attB2 was cut by XhoI, which produced two fragments, one 10.5 kb fragment (pCR2.1-attB1-loxP-y-loxP-frt-attB2 back bone) and one 9 kb fragment* (lacO *repeats) as illustrated by lanes 1, 2 and 4. Lane 6 is the backbone vector pCR2.1-attB1-loxP-y-loxP-frt-attB2. The junction sequence was further verified by sequencing using the FRT R primer. C. The insertion of pCR2.1-attB1-loxP-y-loxP-frt-lacO-attB2 into flies was checked by PCR using attB2 F-white R primers, 1198 F-attB4/09 R, and 1198 F/Y 5' Rev. Lane 1, OR; lane 2,* 1198-lacO *flies. See Methods for experimental details.*

### Verification of *lacO* P element transposition from chromosome 2L to X

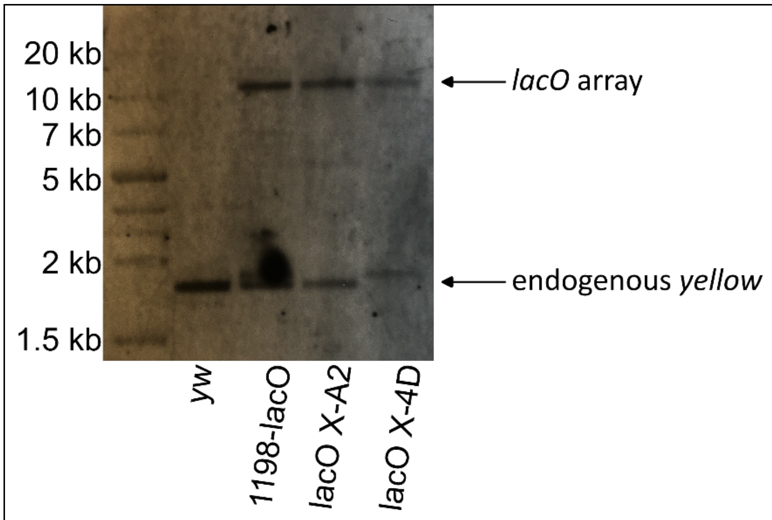


*Fig. S2. Confirmation of an intact lacO repeat cassette after P element mobilization. Genomic DNA from the original 1198* lacO *line, two lines with the* lacO *P element transposed to the X chromosome, and a* yw *control stock with no lacO inserts, was digested with PvuII and probed for DNA sequence of the yellow gene. Probes hybridized to a fragment greater than 10 kb that corresponds to the predicted 11.4 kb fragment containing the* lacO *reporter, and to a 1.5-2 kb fragment corresponding to the endogenous* yellow *gene. See Methods section on lacO transposition mutagenesis screen for experimental details.*

### Assessment of possible suppressors of variegation

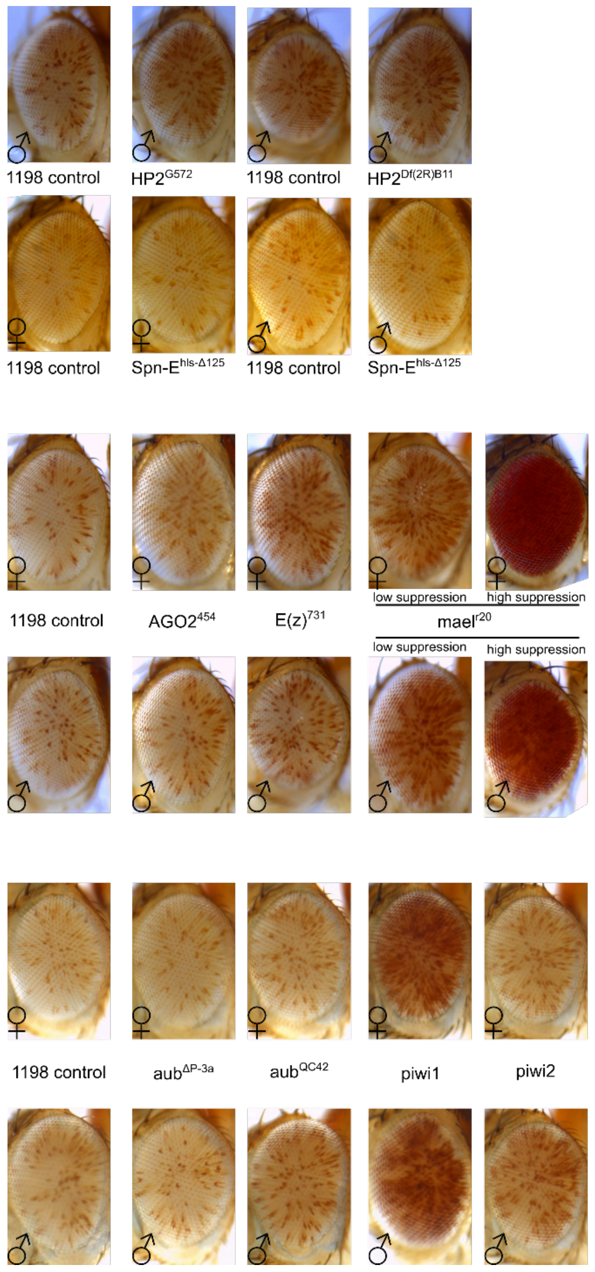


*Figure S3. Assessment of possible suppressors of the* lacO *repeat-induced PEV.* 1198-lacO *flies were crossed to flies carrying genomic mutations for other candidate interacting loci. Variegation in 1198* lacO*/mutant progeny were compared to the following controls: HP2 compared to* cn bw *control;* spn-E *compared to* TM3 Sb *control;* AGO2, E(z) *and* mael *compared to* yw *control;* aub *and* piwi *compared to* yw *control. Suppression of PEV by* piwi1 *is judged likely spurious, given the negative results with* piwi2 *and mutations in* aub, AGO2*, and* Spn-E*.*

### Characterization of new lacO insertion lines

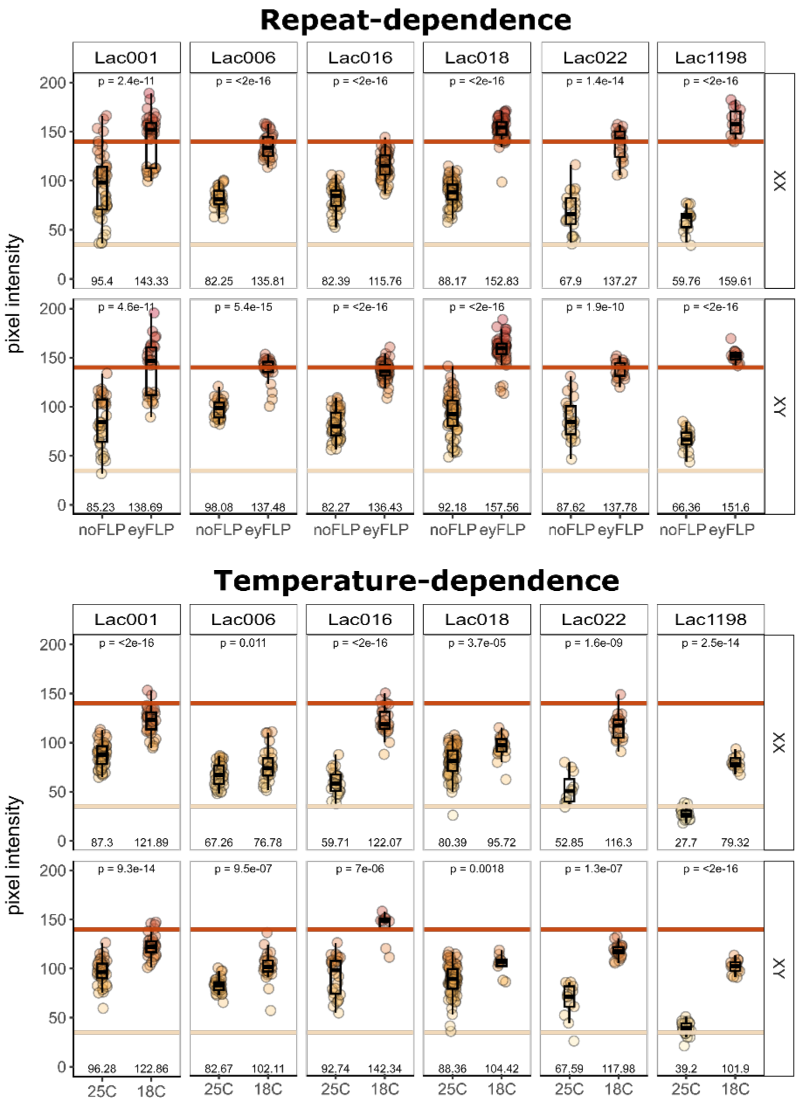


*Figure S4. New* lacO *lines exhibit repeat- and temperature-dependent silencing. Five newly isolated insertional mutants were characterized alongside the ancestral 1198-lacO To test temperature-dependence, yw x lacO crosses were incubated at either 18°C or 25°C before collection of adult progeny from both conditions. To test repeat-dependence lacO males were crossed to females heterozygous for eyFLP expression at 25°C before collecting adult progeny. Flies with lacO and eyFLP were compared to flies with lacO and no FLP. Individual progeny from one or more crosses were frozen and then imaged using a high-throughput macro photography described in (Pipkin et al. 2024). Eye pigment intensity was extracted from photos using ImageJ. Each filled circle represents the average pixel intensity of a single eye; group averages are shown numerically below each experimental group. Box plots show median and interquartile range with whiskers showing 1.5 * IQR. Unadjusted p values for each experimental comparison are shown at the top of each panel. Beige and red horizontal lines denote the average pixel intensity value for* white *and wild-type stocks respectively*

#### Results for HDAC and Sirtuin genes tested by RNAi

**Table S2. TRiP screen for HDACs**

| **TRiP Alleles** | **Screen 1 Impact** | **Screen 2 Impact** |
|---|---|---|
| HDAC3$^{JF01420}$ | Lethal | |
| HDAC3$^{HMS00087}$ | N/A (parental line unhealthy) | |
| HDAC11$^{HMS00483}$ | Su(var) | None |
| Sirt4$^{HMS00944}$ | Lethal | |
| Sirt4$^{GL00548}$ | Lethal | |
| Sirt6$^{JF01583}$ | None | None |
| Sirt6$^{HMS01009}$ | None | None |
| Sirt7$^{JF01558}$ | None | None |
| Sirt7$^{GL01009}$ | E(var) | E(var) |
| Sir2$^{HMJ21708}$ | | Lethal |

### Supplemental Discussion

#### HP1-dependency

The local accumulation of HP1a appears sufficient to trigger variegation; this could occur either through recruitment of additional binding partners or through HP1a-driven phase separation. Earlier studies from the Wallrath lab showed that tethering of HP1a to the *lacO* repeat array used here by an HP1a-lacI fusion protein is sufficient to induce silencing at a wide range of euchromatic insertion sites (Li et al. 2003; Danzer and Wallrath 2004). The appearance of chromatin fibers between the tethering site and distant heterochromatic locations (Li et al. 2003) suggests that the tethered HP1a functions by re-localizing the array to a mass of endogenous constitutive heterochromatin. This could occur through dimerization of HP1a via its chromo shadow domain or through interactions with other HP1a binding proteins, in both cases potentially driving the phase-separation mechanism recently described to explain association of HP1a-containing complexes (Larson et al. 2017; Strom et al. 2017).

HP1a physically interacts with over 100 other proteins and (at least) five RNAs (FlyBase release FB2025_01 (Öztürk-Çolak et al. 2024)). Examining the different heterochromatin forms at different genomic sites, one sees considerable variation in the pattern of HP1a protein partners, assessed by looking for loss of silencing in the presence of a loss-of-function mutation (Phalke et al. 2009). The *lacO* repeat-induced silencing is similar to many other sites in that it is dependent on SU(VAR)3-7 (Figure 2C), a common partner of HP1a that also appears to have a structural role, as shown by an antipodal response to gene dosage (Reuter et al. 1990). In contrast, it is not sensitive to mutations in the gene for HP2 (Table 1), a partner of HP1a with impacts on both heterochromatin formation and mitotic chromosome structure (Shaffer et al. 2006). The functional implications of these choices of partners are unknown. *We conclude that the variegating phenotype observed here is due to local heterochromatin formation dependent on* HP1a *and* SU(VAR)3-7*, but in other ways distinct from other common forms of heterochromatin.*

#### Temperature sensitivity

We observed an unexpectedly strong temperature-dependence for *lacO* repeat-induced silencing, which was enhanced at 25°C and suppressed at 18°C in marked contrast to observations that *white* variegation due to juxtaposition to naturally occurring *D. melanogaster* heterochromatin is suppressed at higher temperatures (Gowen and Gay 1933). Studies of the heterochromatic gene *lt,* which variegates when juxtaposed with euchromatin, indicate that it follows a pattern similar to *lacO* repeat-induced silencing, with elevated temperatures enhancing variegation (first observed in (Gersh 1949) and reviewed in (Girton and Johansen 2008)). These results imply that a key component of heterochromatin formation, or simply maintenance of a stable heterochromatin structure, performs a bit better at lower temperatures, whereas gene expression is a bit better at higher temperatures. These results are consistent with the notion that heterochromatin is a relatively constant chromatin assembly whereas euchromatin is a more dynamic structure, facilitating "unpackaging" to allow transcription. *The unexpected loss of lacO repeat-induced silencing at lower temperatures points to temperature sensitivity in a component uniquely required for this heterochromatin form.*

### Genome defense against repetitive DNA

The small proportion of the *Drosophila* genome that reacts to insertion of the foreign *lacO* DNA repeats by heterochromatin assembly is surprising, given that expression of such repetitious elements is generally deleterious. In other systems silencing of such transgenes can be much more widespread. For example, triplet-repeat induced variegation in mice appears to be independent of the chromosomal location (Saveliev et al. 2003) while in *Drosophila* we find a failure to induce variegation at most sites of insertion. Could there be a failure in recognition, licensing, or maintenance of heterochromatin at these sites? Could heterochromatin be forming over the *lacO* repeats, but failing to spread to the adjacent reporter *w* gene? What can we infer from examination of endogenous repetitive sequences in *Drosophila*?

Endogenous repetitive sequences in eukaryotic genomes broadly fall into two classes: first, the transposable elements and their remnants; second, the tandem arrays, from di- and tri-nucleotides to whole genes. Silencing of both types of elements by heterochromatin packaging is crucial. Silencing prevents transcription and mobilization of the TEs (limiting their impact as mutagens) and stabilizes the genome against spurious recombination and DNA damage triggered by both types of repeats (Janssen et al. 2018). Stable heterochromatin is required to maintain both the short repeats found in satellite DNA and the tandem array of the much larger rRNA genes; loss of HP1a results in excision of circular arrays of these sequences (Peng and Karpen 2007). In our study it is notable that the same small region of the genome is effective in achieving stable heterochromatin formation both for a TE (*1360* or *Invader4*) insert and for the *lacO* tandem repeats. These two types of repetitious DNA appear to be recognized for silencing by different mechanisms.

Across eukaryotic systems transposons are silenced by a variety of RNAi-based mechanisms (Kelleher et al. 2020). In flies, the piRNA system uses captured genomic copies of TE sequences to silence active TEs – a kind of genetic acquired immunity (Czech et al. 2018) -- and this recognition system is used to silence our P-element constructs carrying *1360* or *Invader4* at the 1198 site (Sentmanat and Elgin 2012). Invasion of *D. melanogaster* by a novel TE invokes a rapid response to incorporate the new TE into the piRNA silencing system (Ho et al. 2024; Khurana et al. 2011). However, the piRNA system does not appear to be directly applicable for the *lacO* repeats, as we found no endogenous *D. melanogaster* genome DNA sequences with significant homology to the *lacO* sequence, and our genetic tests found no sensitivity to mutations in components of the piRNA pathway (Table 1, Figure S3). We conclude that the feature that defines the ~1.5% of the *Drosophila melanogaster* euchromatic genome as the portion that supports ectopic heterochromatin formation in response to inserted repetitious sequences is not concerned with the "recognition" step but could be critical for licensing or maintenance of heterochromatin. Alternatively, there could be a failure in spreading of heterochromatin to the *white* reporter gene. The *lacO* repeat-induced silencing may be particularly vulnerable, in that it does not appear to utilize H3K9me2/3, a common element used (but not always required) for spreading (Danzer and Wallrath 2004; Hines et al. 2009). However, heterochromatin formation at site 1198 induced by a TE insertion does utilize H3K9me2/3 (Sentmanat and Elgin 2012), and this construct is similarly found to induce heterochromatin formation at the same limited set of genomic sites, making this explanation unlikely. *The question remains unresolved.*

### Recognition of novel repetitive elements

How might a tandem array of a novel foreign DNA sequence (here the *E. coli lacO* fragment) be recognized as deleterious and silenced? We are hampered by the lack of an established mechanism for recognition of the tandem repeats as a class. In fact, it appears that several different mechanisms are in play to recognize endogenous tandem repeats, triggering silencing in each case. In the broadest case, it has been demonstrated that differences in GC content of inserted foreign DNA can have an impact on heterochromatin vs euchromatin packaging in *S. cerevisiae;* transgenic bacterial DNA with low GC content is favored for heterochromatin packaging (Meneu et al. 2025). This bias might also impact the packaging of tandem arrays. While the endogenous *D. melanogaster* satellite DNAs are often AT-rich, the GC content of the *lacO* fragment used here is 44%, similar to the *D. melanogaster* genome as a whole, eliminating this characteristic of DNA as a potential recognition feature.

Focusing on *Drosophila*, one notes that other structural features of DNA present possibilities. For example, some AT-rich satellite DNA sequences bind the D1 protein (with 10 AT-hook domains) through the minor groove (Aulner et al. 2002; Blattes et al. 2006). Interestingly, while D1 interacts with HP1a, it also interacts with components of the transposon-silencing piRNA pathway (Chavan et al. 2025). Other satellite DNAs specifically bind to known chromosomal proteins (e.g., GAF binds AAGAG repeats (Gaskill et al. 2023)) or other maternally provided proteins, as yet uncharacterized (Yuan and O'Farrell 2016). In the two major instances in *D. melanogaster* where genes are found in a tandem array, the rRNA genes (80-600 copies, (Lu et al. 2018)) and the histone genes (~110 copies (Crain et al. 2024)), the loci are identified for heterochromatin packaging through specific protein binding (NoRC for the rRNA genes, (Guetg et al. 2010); HERS for the histone genes, (Ito et al. 2012)). These diverse mechanisms all drive heterochromatin formation through localization of appropriate histone modifying enzymes and heterochromatin structural proteins. *However, these strategies do not appear to be well-suited for dealing with novel invading DNA repeats, as they are not applicable beyond a specific subset of cases (e.g., short AT-rich tandem repeats). There is no reason to anticipate the presence of a eukaryotic nuclear protein that can specifically recognize the lacO repeat as a foreign invader to be silenced.*

Evidence from multiple systems implicates pausing by RNA polymerase (e.g., *Homo sapiens* (Groh et al. 2014); *Caenorhabditis elegans*, (Zeller et al. 2016)) or DNA polymerase (Gaggioli et al. 2023), leading to formation of R-loops and/or D-loops at repetitive sites as a trigger for silencing. In particular, epigenetic silencing of the Fragile X locus (CGG repeats) is dependent on promoter-bound trinucleotide repeat mRNA (Colak et al. 2014; Loomis et al. 2014). Similar results have been obtained for the Friedreich's Ataxia locus (GAA repeats) (Groh et al. 2014). These results argue in favor of direct recognition of the stalled transcription complex and/or the resulting RNA/DNA hybrid, rather than a model that invokes RNAi (Colak et al. 2014). Studies of $\alpha$–satellite repeats in mammals have suggested that such RNA-DNA hybrids can form a platform for direct stable binding of H3K9 HMTs, particularly SUV39H1, which can drive heterochromatin assembly (reviewed by (Stamidis and Żylicz 2023)). However, the HMTs do not seem to be involved in the silencing of the *lacO*-repeats. Alternatively, HP1a is similar in having a capacity to bind to nucleic acids directly, in addition to its interaction with H3K9me2/3 and with numerous other chromosomal proteins (Zhao et al. 2000; Muchardt et al. 2002). Thus it has been suggested that heterochromatin might form at an R-loop through direct binding of any of several heterochromatin proteins. Further, the R-loop can lead to a stalled replication fork, resulting in D-loops, DNA damage and repair. The complexes created for DNA repair also localize heterochromatin proteins to the site, notably HP1a and HDAC1/2 (e.g., (Luijsterburg et al. 2009; Miller et al. 2010; Zeller and Gasser 2017), reviewed in (Lemaître and Soutoglou 2014)). Localization of HP1a might also be enhanced by its interactions with radiation repair scaffolding protein MU2 (Dronamraju and Mason 2011) and with NBS1 (Nijmegen Breakage Syndrome 1), part of the MRN complex involved in resection of damaged DNA and complex docking (Bosso et al. 2019). Localization of HP1a and HDAC1/2 might be sufficient to drive a repetitious site into a heterochromatic membrane-less compartment, stabilizing heterochromatin formation. *Such a mechanism would be broadly applicable in that the actual DNA sequence of the repeat appears to be of no consequence; the critical feature for R-loop and/or D-loop formation is the tandem array of repeats. As pointed out by Zeller & Gasser* (Zeller and Gasser 2017)*, this mechanism is well suited for recognizing insertions of foreign DNA (if tandem repeats) as sites to be silenced.*

Using the above model, what features of the repeat array might limit successful formation of ectopic heterochromatin? Mechanistically, a major factor in generating stable heterochromatin in response to the inserted repeats could be the local concentration of HP1a, with the limiting factor being the requirement for a sufficient concentration of that protein to draw that DNA region into the HP1a-rich droplets (membraneless particles) resulting from phase separation (Strom et al. 2017; Larson et al. 2017). Modeling to reproduce Hi-C data suggests that 20 kb or more is required to generate a stable heterochromatin phase (MacPherson et al. 2018), while our *lacO* array is 9.2 kb. However, significantly smaller arrays (in total nts) of tandem triplet repeats do generate ectopic heterochromatin in humans, causing silencing of the adjacent gene in Fragile X syndrome (~200-600 nts CGG needed) and Friedreich's Ataxia (40-500 nts GAA needed) (Saveliev et al. 2003; Warren and Nelson 1994). Reviewing the data available, we do not see any general relationship between susceptibility to silencing and the total length of the repetitious array.

Could the number of copies in a tandem array, rather than the size of the individual repeated element or the final size of the tandem array, be the critical parameter? For example, there are many sites with small numbers of tandem repeats of CGG and GAA scattered throughout the euchromatin (Katti et al. 2001; Astolfi et al. 2003), and these are not reported to drive heterochromatin formation. While we have not seen studies that address the question in general, the human disease sites are informative. Specifically, arrays with fewer than 200 copies of the CGG triplet found in Fragile X do not drive heterochromatin formation, while a higher copy number does (Warren and Nelson 1994). Similarly, there are many examples from a variety or organisms of tandem gene duplications within euchromatic regions where the genes are appropriately expressed, providing more of a desired product (e.g., (Perry et al. 2007; Bouchebti et al. 2024; Klure et al. 2025)) or allowing for the emergence of isoforms (Grönke et al. 2010); both outcomes are observed as part of the evolutionary process. These cases usually have ca. two to ten copies of the gene in question, suggesting a higher threshold for the number of copies required to trigger heterochromatin formation. Indeed, while the ~110 copies of the *D. melanogaster* histone genes appear to be packaged as heterochromatin (Ner et al. 2002; Huisinga et al. 2016; van Steensel et al. 2001), the minimal construct with 12 copies at the same genomic site does not appear to be packaged as heterochromatin (Ahmad et al. 2025), even though both arrays generate the HLB (histone locus body) structure. The significance of copy number is in congruence with models based on the use of R-loops or D-loops to trigger heterochromatin formation. Studies aimed at finding a direct molecular link between R-loops and the pathology of triplet repeats have found that increasing R-loop levels (using camptothecin) leads to increased levels of repressive chromatin marks at the triplet repeat locus, and that the level of R-loops correlates with expansion length (number of triplet repeats) (Groh et al. 2014). *Considering all of the available evidence, a strategy based on generation of R-loops and/or D-loops seems the most likely mechanism for the immediate recognition of the* lacO *tandem repeats for the silencing seen in* **Figure 1**. *The number of tandem repeats in an array appears to be the critical parameter for achieving silencing.*

### Formation of ectopic heterochromatin

As discussed above, a likely model for recognition of the *lacO*-repeat array as a target for silencing by heterochromatin formation is RNA- and/or DNA-loop formation. Several reports demonstrate that formation of the ensuing DNA damage repair complex involves recruitment of HP1 and HDACs to the site. These indeed are the heterochromatin components that we find to be critical for reporter silencing proximal to the *lacO* repeats. While the presence of HP1a could promote heterochromatin formation by promoting association with the HP1a-enriched membraneless domains, silencing could also be created by the increase in nucleosome occupancy postulated on histone deacetylation. It has also been suggested that deacetylation of histones prevents the engagement of chromatin remodeling enzymes, that bind to the acetylated histone modifications via their bromodomain and destabilize nucleosomes to promote transcription (Swygert and Peterson 2014). This supports the hypothesis that in this case, the process of histone deacetylation and presence of HP1a are sufficient for licensing and maintenance of the heterochromatin state. *Further investigation of this lacO repeat-induced silencing, and of the repeat arrays that can trigger it, would no doubt be informative.*

If RNA- and/or DNA-loop formation are required for *lacO* repeat-induced silencing, the lacO-repeat insertion sites that support heterochromatin packaging must allow accessibility for transcription, but be amenable to stochastic silencing to produce a variegating phenotype. A documented example of such a sensitive system is the agouti mouse, where insertion of an IAP (TE Intracisternal A-particle) results in variable coat color (and associated traits) within a litter of pups based on variable ectopic expression of a downstream gene. This case reflects metastable DNA methylation (5'mC) of the IAP transposon (Bertozzi and Ferguson-Smith 2020). While a screen identified ~100 such IAP insertions across the genome, only in rare instances did metastable elements act as promoters impacting a nearby gene (Kazachenka et al. 2018). While the details will differ in flies (which have little or no persistent 5'mC), the concept that insertion of a given repetitious element will produce a visible phenotype at only a few sites in the genome could be general. *The process that produces a metastable chromatin modification, and determines its capacity to impact adjacent genes, remains to be explored. It should be possible to exploit the system developed here for that purpose.*

Astolfi, Paola, Dina Bellizzi, and Vittorio Sgaramella. 2003. “Frequency and Coverage of Trinucleotide Repeats in Eukaryotes.” *Gene*, Papers presented at the meeting “Molecular

Evolution: evolution, genomics, bioinformatics" (Sorrento, June 13-16, 2002) - Part 1, vol. 317 (October): 117–25. https://doi.org/10.1016/S0378-1119(03)00659-0.

Aulner, Nathalie, Monod ,Caroline, Mandicourt ,Guillaume, et al. 2002. "The AT-Hook Protein D1 Is Essential for Drosophila Melanogaster Development and Is Implicated in Position-Effect Variegation." *Molecular and Cellular Biology* 22 (4): 1218–32. https://doi.org/10.1128/MCB.22.4.1218-1232.2002.

Bertozzi, Tessa M., and Anne C. Ferguson-Smith. 2020. "Metastable Epialleles and Their Contribution to Epigenetic Inheritance in Mammals." *Seminars in Cell & Developmental Biology*, SI: Chromatin dynamics in regeneration, vol. 97 (January): 93–105. https://doi.org/10.1016/j.semcdb.2019.08.002.

Blattes, Roxane, Caroline Monod, Guillaume Susbielle, et al. 2006. "Displacement of D1, HP1 and Topoisomerase II from Satellite Heterochromatin by a Specific Polyamide." *The EMBO Journal* 25 (11): 2397–408. https://doi.org/10.1038/sj.emboj.7601125.

Bosso, Giuseppe, Francesca Cipressa, Maria Lina Moroni, et al. 2019. "NBS1 Interacts with HP1 to Ensure Genome Integrity." *Cell Death & Disease* 10 (12): 1–15. https://doi.org/10.1038/s41419-019-2185-x.

Bouchebti, Sofia, Yael Gershon, Alexander Gordin, Dorothée Huchon, and Eran Levin. 2024. "Tolerance and Efficient Metabolization of Extremely High Ethanol Concentrations by a Social Wasp." *Proceedings of the National Academy of Sciences* 121 (44): e2410874121. https://doi.org/10.1073/pnas.2410874121.

Chavan, Ankita, Lena Skrutl, Federico Uliana, et al. 2025. "Multi-Tissue Characterization of the Constitutive Heterochromatin Proteome in Drosophila Identifies a Link between Satellite DNA Organization and Transposon Repression." *PLOS Biology* 23 (1): e3002984. https://doi.org/10.1371/journal.pbio.3002984.

Colak, Dilek, Nikica Zaninovic, Michael S. Cohen, et al. 2014. "Promoter-Bound Trinucleotide Repeat mRNA Drives Epigenetic Silencing in Fragile X Syndrome." *Science* 343 (6174): 1002–5. https://doi.org/10.1126/science.1245831.

Crain, Aaron T., Megan B. Butler, Christina A. Hill, Mai Huynh, Robert K. McGinty, and Robert J. Duronio. 2024. "Drosophila Melanogaster Set8 and L(3)Mbt Function in Gene Expression Independently of Histone H4 Lysine 20 Methylation." *Genes & Development* 38 (9–10): 455–72. https://doi.org/10.1101/gad.351698.124.

Czech, Benjamin, Marzia Munafò, Filippo Ciabrelli, et al. 2018. "piRNA-Guided Genome Defense: From Biogenesis to Silencing." *Annual Review of Genetics* 52 (November): 131–57. https://doi.org/10.1146/annurev-genet-120417-031441.

Danzer, John R., and Lori L. Wallrath. 2004. "Mechanisms of HP1-Mediated Gene Silencing in Drosophila." *Development* 131 (15): 3571–80. https://doi.org/10.1242/dev.01223.

Dronamraju, Raghuvar, and James M. Mason. 2011. "MU2 and HP1a Regulate the Recognition of Double Strand Breaks in Drosophila Melanogaster." *PLOS ONE* 6 (9): e25439. https://doi.org/10.1371/journal.pone.0025439.

Gaggioli, Vincent, Calvin S. Y. Lo, Nazaret Reverón-Gómez, et al. 2023. “Dynamic de Novo Heterochromatin Assembly and Disassembly at Replication Forks Ensures Fork Stability.” *Nature Cell Biology* 25 (7): 1017–32. https://doi.org/10.1038/s41556-023-01167-z.

Gaskill, Marissa M., Isabella V. Soluri, Annemarie E. Branks, et al. 2023. “Localization of the Drosophila Pioneer Factor GAF to Subnuclear Foci Is Driven by DNA Binding and Required to Silence Satellite Repeat Expression.” *Developmental Cell* 58 (17): 1610-1624.e8. https://doi.org/10.1016/j.devcel.2023.06.010.

Gersh, E. S. 1949. “Influence of Temperature on the Expression of Position Effects in the Scute-8 Stock of Drosophila Melanogaster and Its Relation to Heterochromatization.” *Genetics* 34 (6): 701–7. https://doi.org/10.1093/genetics/34.6.701.

Girton, Jack R., and Kristen M. Johansen. 2008. “Chromatin Structure and the Regulation of Gene Expression: The Lessons of PEV in Drosophila.” In *Long-Range Control of Gene Expression*, vol. 61. Advances in Genetics. Elsevier. https://doi.org/10.1016/S0065-2660(07)00001-6.

Gowen, John W., and E. H. Gay. 1933. “Effect of Temperature on Eversporting Eye Color in Drosophila Melanogaster.” *Science* 77 (1995): 312–312. https://doi.org/10.1126/science.77.1995.312.a.

Groh, Matthias, Michele M. P. Lufino, Richard Wade-Martins, and Natalia Gromak. 2014. “R-Loops Associated with Triplet Repeat Expansions Promote Gene Silencing in Friedreich Ataxia and Fragile X Syndrome.” *PLoS Genetics* 10 (5): e1004318. https://doi.org/10.1371/journal.pgen.1004318.

Grönke, Sebastian, David-Francis Clarke, Susan Broughton, T. Daniel Andrews, and Linda Partridge. 2010. “Molecular Evolution and Functional Characterization of Drosophila Insulin-Like Peptides.” *PLOS Genetics* 6 (2): e1000857. https://doi.org/10.1371/journal.pgen.1000857.

Guetg, Claudio, Philipp Lienemann, Valentina Sirri, et al. 2010. “The NoRC Complex Mediates the Heterochromatin Formation and Stability of Silent rRNA Genes and Centromeric Repeats.” *The EMBO Journal* 29 (13): 2135–46. https://doi.org/10.1038/emboj.2010.17.

Hines, Karrie A, Diane E Cryderman, Kaitlin M Flannery, et al. 2009. “Domains of Heterochromatin Protein 1 Required for Drosophila Melanogaster Heterochromatin Spreading.” *Genetics* 182 (4): 967–77. https://doi.org/10.1534/genetics.109.105338.

Ho, Samantha, William Theurkauf, and Nicholas Rice. 2024. “piRNA-Guided Transposon Silencing and Response to Stress in Drosophila Germline.” *Viruses* 16 (5): 5. https://doi.org/10.3390/v16050714.

Huisinga, Kathryn L., Nicole C. Riddle, Wilson Leung, et al. 2016. “Targeting of P-Element Reporters to Heterochromatic Domains by Transposable Element 1360 in Drosophila Melanogaster.” *Genetics* 202 (2): 565–82. https://doi.org/10.1534/genetics.115.183228.

Ito, Saya, Sally Fujiyama-Nakamura, Shuhei Kimura, et al. 2012. “Epigenetic Silencing of Core Histone Genes by HERS in Drosophila.” *Molecular Cell* 45 (4): 494–504. https://doi.org/10.1016/j.molcel.2011.12.029.

Janssen, Aniek, Serafin U. Colmenares, and Gary H. Karpen. 2018. “Heterochromatin: Guardian of the Genome.” *Annual Review of Cell and Developmental Biology* 34 (1): 265–88. https://doi.org/10.1146/annurev-cellbio-100617-062653.

Katti, Mukund V., Prabhakar K. Ranjekar, and Vidya S. Gupta. 2001. “Differential Distribution of Simple Sequence Repeats in Eukaryotic Genome Sequences.” *Molecular Biology and Evolution* 18 (7): 1161–67. https://doi.org/10.1093/oxfordjournals.molbev.a003903.

Kazachenka, Anastasiya, Tessa M. Bertozzi, Marcela K. Sjoberg-Herrera, et al. 2018. “Identification, Characterization, and Heritability of Murine Metastable Epialleles: Implications for Non-Genetic Inheritance.” *Cell* 175 (5): 1259-1271.e13. https://doi.org/10.1016/j.cell.2018.09.043.

Kelleher, Erin S., Daniel A. Barbash, and Justin P. Blumenstiel. 2020. “Taming the Turmoil Within: New Insights on the Containment of Transposable Elements.” *Trends in Genetics* 36 (7): 474–89. https://doi.org/10.1016/j.tig.2020.04.007.

Khurana, Jaspreet S., Jie Wang, Jia Xu, et al. 2011. “Adaptation to P Element Transposon Invasion in Drosophila Melanogaster.” *Cell* 147 (7): 1551–63. https://doi.org/10.1016/j.cell.2011.11.042.

Klure, Dylan M., Robert Greenhalgh, Teri J. Orr, Michael D. Shapiro, and M. Denise Dearing. 2025. “Parallel Gene Expansions Drive Rapid Dietary Adaptation in Herbivorous Woodrats.” *Science* 387 (6730): 156–62. https://doi.org/10.1126/science.adp7978.

Larson, Adam G., Daniel Elnatan, Madeline M. Keenen, et al. 2017. “Liquid Droplet Formation by HP1α Suggests a Role for Phase Separation in Heterochromatin.” *Nature* 547 (7662): 236–40. https://doi.org/10.1038/nature22822.

Lemaître, Charlène, and Evi Soutoglou. 2014. “Double Strand Break (DSB) Repair in Heterochromatin and Heterochromatin Proteins in DSB Repair.” *DNA Repair*, Cutting-edge Perspectives in Genomic Maintenance, vol. 19 (July): 163–68. https://doi.org/10.1016/j.dnarep.2014.03.015.

Li, Yuhong, John R. Danzer, Pedro Alvarez, Andrew S. Belmont, and Lori L. Wallrath. 2003. “Effects of Tethering HP1 to Euchromatic Regions of the Drosophila Genome.” *Development (Cambridge, England)* 130 (9): 1817–24.

Loomis, Erick W., Lionel A. Sanz, Frédéric Chédin, and Paul J. Hagerman. 2014. “Transcription-Associated R-Loop Formation across the Human FMR1 CGG-Repeat Region.” *PLOS Genetics* 10 (4): e1004294. https://doi.org/10.1371/journal.pgen.1004294.

Lu, Kevin L, Jonathan O Nelson, George J Watase, Natalie Warsinger-Pepe, and Yukiko M Yamashita. 2018. “Transgenerational Dynamics of rDNA Copy Number in Drosophila Male Germline Stem Cells.” *eLife* 7 (February): e32421. https://doi.org/10.7554/eLife.32421.

Luijsterburg, Martijn S., Christoffel Dinant, Hannes Lans, et al. 2009. “Heterochromatin Protein 1 Is Recruited to Various Types of DNA Damage.” *Journal of Cell Biology* 185 (4): 577–86. https://doi.org/10.1083/jcb.200810035.

MacPherson, Quinn, Bruno Beltran, and Andrew J. Spakowitz. 2018. “Bottom-up Modeling of Chromatin Segregation Due to Epigenetic Modifications.” *Proceedings of the National Academy* *of Sciences of the United States of America* 115 (50): 12739–44. https://doi.org/10.1073/pnas.1812268115.

Meneu, Léa, Christophe Chapard, Jacques Serizay, et al. 2025. “Sequence-Dependent Activity and Compartmentalization of Foreign DNA in a Eukaryotic Nucleus.” *Science (New York, N.Y.)* 387 (6734): eadm9466. https://doi.org/10.1126/science.adm9466.

Miller, Kyle M., Jorrit V. Tjeertes, Julia Coates, et al. 2010. “Human HDAC1 and HDAC2 Function in the DNA-Damage Response to Promote DNA Nonhomologous End-Joining.” *Nature Structural & Molecular Biology* 17 (9): 1144–51. https://doi.org/10.1038/nsmb.1899.

Muchardt, Christian, Marie Guillemé, Jacob-S Seeler, Didier Trouche, Anne Dejean, and Moshe Yaniv. 2002. “Coordinated Methyl and RNA Binding Is Required for Heterochromatin Localization of Mammalian HP1α.” *EMBO Reports* 3 (10): 975–81. https://doi.org/10.1093/embo-reports/kvf194.

Ner, Sarbjit S, Michael J Harrington, and Thomas A Grigliatti. 2002. “A Role for the Drosophila SU(VAR)3-9 Protein in Chromatin Organization at the Histone Gene Cluster and in Suppression of Position-Effect Variegation.” *Genetics* 162 (4): 1763–74. https://doi.org/10.1093/genetics/162.4.1763.

Öztürk-Çolak, Arzu, Steven J Marygold, Giulia Antonazzo, et al. 2024. “FlyBase: Updates to the Drosophila Genes and Genomes Database.” *Genetics* 227 (1): iyad211. https://doi.org/10.1093/genetics/iyad211.

Peng, Jamy C., and Gary H. Karpen. 2007. “H3K9 Methylation and RNA Interference Regulate Nucleolar Organization and Repeated DNA Stability.” *Nature Cell Biology* 9 (1): 25–35. https://doi.org/10.1038/ncb1514.

Perry, George H., Nathaniel J. Dominy, Katrina G. Claw, et al. 2007. “Diet and the Evolution of Human Amylase Gene Copy Number Variation.” *Nature Genetics* 39 (10): 1256–60. https://doi.org/10.1038/ng2123.

Phalke, Sameer, Olaf Nickel, Diana Walluscheck, Frank Hortig, Maria Cristina Onorati, and Gunter Reuter. 2009. “Retrotransposon Silencing and Telomere Integrity in Somatic Cells of Drosophila Depends on the Cytosine-5 Methyltransferase DNMT2.” *Nature Genetics* 41 (6): 696–702. https://doi.org/10.1038/ng.360.

Pipkin, Heidi J. J., Hunter L. Lindsay, Adam T. Smiley, Jack D. Jurmu, and Andrew M. Arsham. 2024. “An Accessible Digital Imaging Workflow for Multiplexed Quantitative Analysis of Adult Eye Phenotypes in Drosophila Melanogaster.” Preprint, bioRxiv, August 28. https://doi.org/10.1101/2024.01.26.577286.

Reuter, G., M. Giarre, J. Farah, J. Gausz, A. Spierer, and P. Spierer. 1990. “Dependence of Position-Effect Variegation in Drosophila on Dose of a Gene Encoding an Unusual Zinc-Finger Protein.” *Nature* 344 (6263): 219–23. https://doi.org/10.1038/344219a0.

Saveliev, Alexander, Christopher Everett, Tammy Sharpe, Zoë Webster, and Richard Festenstein. 2003. “DNA Triplet Repeats Mediate Heterochromatin-Protein-1-Sensitive Variegated Gene Silencing.” *Nature* 422 (6934): 909–13. https://doi.org/10.1038/nature01596.

Sentmanat, Monica F., and Sarah C. R. Elgin. 2012. “Ectopic Assembly of Heterochromatin in Drosophila Melanogaster Triggered by Transposable Elements.” *Proceedings of the National Academy of Sciences of the United States of America* 109 (35): 14104–9. https://doi.org/10.1073/pnas.1207036109.

Shaffer, Christopher D, Giovanni Cenci, Brandi Thompson, et al. 2006. “The Large Isoform of Drosophila Melanogaster Heterochromatin Protein 2 Plays a Critical Role in Gene Silencing and Chromosome Structure.” *Genetics* 174 (3): 1189–204. https://doi.org/10.1534/genetics.106.057604.

Stamidis, Nikolaos, and Jan Jakub Żylicz. 2023. “RNA-mediated Heterochromatin Formation at Repetitive Elements in Mammals.” *The EMBO Journal* 42 (8): e111717. https://doi.org/10.15252/embj.2022111717.

Steensel, Bas van, Jeffrey Delrow, and Steven Henikoff. 2001. “Chromatin Profiling Using Targeted DNA Adenine Methyltransferase.” *Nature Genetics* 27 (3): 304–8. https://doi.org/10.1038/85871.

Strom, Amy R., Alexander V. Emelyanov, Mustafa Mir, Dmitry V. Fyodorov, Xavier Darzacq, and Gary H. Karpen. 2017. “Phase Separation Drives Heterochromatin Domain Formation.” *Nature* 547 (7662): 241–45. https://doi.org/10.1038/nature22989.

Swygert, Sarah G., and Craig L. Peterson. 2014. “Chromatin Dynamics: Interplay between Remodeling Enzymes and Histone Modifications.” *Biochimica et Biophysica Acta (BBA) - Gene Regulatory Mechanisms*, Molecular mechanisms of histone modification function, vol. 1839 (8): 728–36. https://doi.org/10.1016/j.bbagrm.2014.02.013.

Warren, Stephen T., and David L. Nelson. 1994. “Advances in Molecular Analysis of Fragile X Syndrome.” *JAMA* 271 (7): 536–42. https://doi.org/10.1001/jama.1994.03510310066040.

Yuan, Kai, and Patrick H. O’Farrell. 2016. “TALE-Light Imaging Reveals Maternally Guided, H3K9me2/3-Independent Emergence of Functional Heterochromatin in Drosophila Embryos.” *Genes & Development* 30 (5): 579–93. https://doi.org/10.1101/gad.272237.115.

Zeller, Peter, and Susan M. Gasser. 2017. “The Importance of Satellite Sequence Repression for Genome Stability.” *Cold Spring Harbor Symposia on Quantitative Biology* 82: 15–24. https://doi.org/10.1101/sqb.2017.82.033662.

Zeller, Peter, Jan Padeken, Robin Van Schendel, Veronique Kalck, Marcel Tijsterman, and Susan M Gasser. 2016. “Histone H3K9 Methylation Is Dispensable for Caenorhabditis Elegans Development but Suppresses RNA:DNA Hybrid-Associated Repeat Instability.” *Nature Genetics* 48 (11): 1385–95. https://doi.org/10.1038/ng.3672.

Zhao, Tao, Thomas Heyduk, C. David Allis, and Joel C. Eissenberg. 2000. “Heterochromatin Protein 1 Binds to Nucleosomes and DNA in Vitro *.” *Journal of Biological Chemistry* 275 (36): 28332–38. https://doi.org/10.1074/jbc.M003493200.
